# Phenotypic dynamics and behaviors of neutrophils during barrier challenge

**DOI:** 10.64898/2026.01.06.697946

**Authors:** Marie Siwicki, Hanjoo Brian Shim, Michelle Newton, Woo-Yong Lee, Paul Kubes

## Abstract

Neutrophils are intrinsically antimicrobial innate immune cells, but they have recently been observed to play diverse roles across various conditions. The concept of neutrophil heterogeneity has thus emerged, yet a fundamental understanding of the mechanisms and processes underlying neutrophil diversity *in vivo* is lacking. Here, we interrogated neutrophils’ native propensities to diversify by studying them in the setting of host defense to barrier challenge. Using both high-parameter, high throughput methods and intravital microscopy, we tracked cellular dynamics, identity, and behaviors and asked how these aligned with critical host defensive functions. We found that neutrophil diversity was underscored by both turnover and plasticity of cells within an infection site, and that environment-specific adaptation occurred rapidly upon extravasation and swarm initiation. The population-level phenotypic landscape shifted considerably through phases of inflammation initiation, peak, and resolution; and adaptation varied substantially between infections and aseptic wounds. Like mature neutrophils, immature neutrophils were capable of phenotypic adaptation in infection sites but were deficient in host defense functions. We unexpectedly identified a novel population of CD101-low, mature neutrophils, expressing elevated levels of PD-L1 and ICAM-1, that was associated with key antimicrobial effector functions and most pronounced during the neutrophil response to large or recalcitrant challenge. Our findings highlight that neutrophil specification is rapid and environment-dependent, and that specific phenotypes are linked to critical host defensive behaviors and functions *in vivo*.

## Introduction

Neutrophils are powerful anti-microbial granulocytes of the innate immune system. In recent years, they have been observed to play important roles in non-infectious contexts as well, including in wound healing, cardiac stress, autoimmunity, and cancer(*1–5*). In several settings, the concept has emerged that neutrophils may be heterogeneous, and neutrophil polarization states or subsets have been highlighted, especially in contexts of cancer and cardiovascular disease(*6, 7*). However, the underlying basis of neutrophil heterogeneity remains largely unknown. While several studies have drawn associations between neutrophil states and factors including resource scarcity and maturational age(*8–12*), the majority of these studies have been in contexts of cancer or other lifestyle diseases—arenas largely departed from the evolutionary pressures that have endowed neutrophils with the capacity to diversify. For this reason, we sought to study the intricacies of neutrophil heterogeneity in an evolutionarily powerful setting: infection and acute injury to a barrier site.

The skin is nominally one of the most important immune organs, protecting the host against invasive pathogens in the external environment. When the skin is compromised, neutrophils are some of the first cellular responders, mobilizing rapidly to stem infection risk via anti-microbial effector functions(*1*). Neutropenia is a major risk factor for recurrent infections and death(*13–15*), highlighting the evolutionarily powerful role these cells play in settings of barrier challenges. Here, we use this setting—infection and wounding of the skin—to establish a foundational picture of neutrophils’ native propensities for diversity, and to ask how this aligns with critical neutrophil behaviors and functions *in vivo*.

A major challenge to studying neutrophil functional heterogeneity has been the quest to use transcriptomics to capture neutrophil subsets or states. Since neutrophils are less transcriptionally active than most other immune cells, they are frequently lost in single cell RNA-seq pipelines(*6, 16*). Furthermore, high levels of intracellular nucleases make it challenging to recover the transcripts that they do express(*17*). In addition, since neutrophils execute a great deal of their protein synthesis during maturation within the bone marrow, transcripts recovered in the periphery may not accurately reflect their protein expression or functionality, best evidenced by low or no transcript for the most abundant protein neutrophil elastase(*18*). Finally, most transcriptional approaches capture only a snapshot of neutrophil heterogeneity, failing to address dynamics of neutrophil diversity within often rapidly changing inflammatory environments.

To circumvent these challenges, we have employed spectral flow cytometry to capture neutrophil heterogeneity on a protein level, enabling high-throughput studies of neutrophil diversity using a core panel of over 20 phenotypic markers. In combination with intravital microscopy and transgenic mouse models that enable tracking neutrophils over time, we have been able to assess neutrophil dynamics and plasticity, as well as relationships between environment, phenotype, and function, during acute barrier challenge.

## Results

### Accumulation and turnover support a scaled neutrophil response during infection

In order to model an evolutionarily relevant infectious challenge to the skin, we employed a slight modification of a previously published model wherein a low infectious dose of *Staphylococcus aureus* (*S. aureus*) is associated with agarose microbeads of 40-100 um diameter. The agarose, functioning as a foreign body, enables the *S. aureus* to form a biofilm, allowing development of a persistent (up to 7+ days) infection with a low bacterial MOI(*19*). Using this approach, we inoculated murine skin with ∼3×10^3^ CFU *S. aureus* and analyzed neutrophil presence in the infection over the course of several days [Fig. 1A-C]. Levels of neutrophils peaked around 24 hours after infection [Fig. 1B], and waned over the course of 7-10 days, with numbers of neutrophils proportional to the burden of *S. aureus* in the skin [Fig. 1C]. Intravital microscopy during the first 24h of infection revealed neutrophils arresting within and exiting from nearby blood vessels, then trafficking toward the site of infection to surround the *S. aureus* microbeads [Fig. 1D]. By 24h, neutrophils were observed in a robust swarm around beads containing GFP expressing *S. aureus*, and live imaging revealed *S. aureus* biofilms being dismantled by neutrophils within the swarm [Fig. 1E]. We noted that a minority of neutrophils appeared to be positive for *S. aureus* by both imaging and flow cytometry, indicating that the magnitude of the neutrophil swarm outscales the requirement for individual cells binding and phagocytosing pathogen [Fig. 1E, S1A].

**Figure 1.**
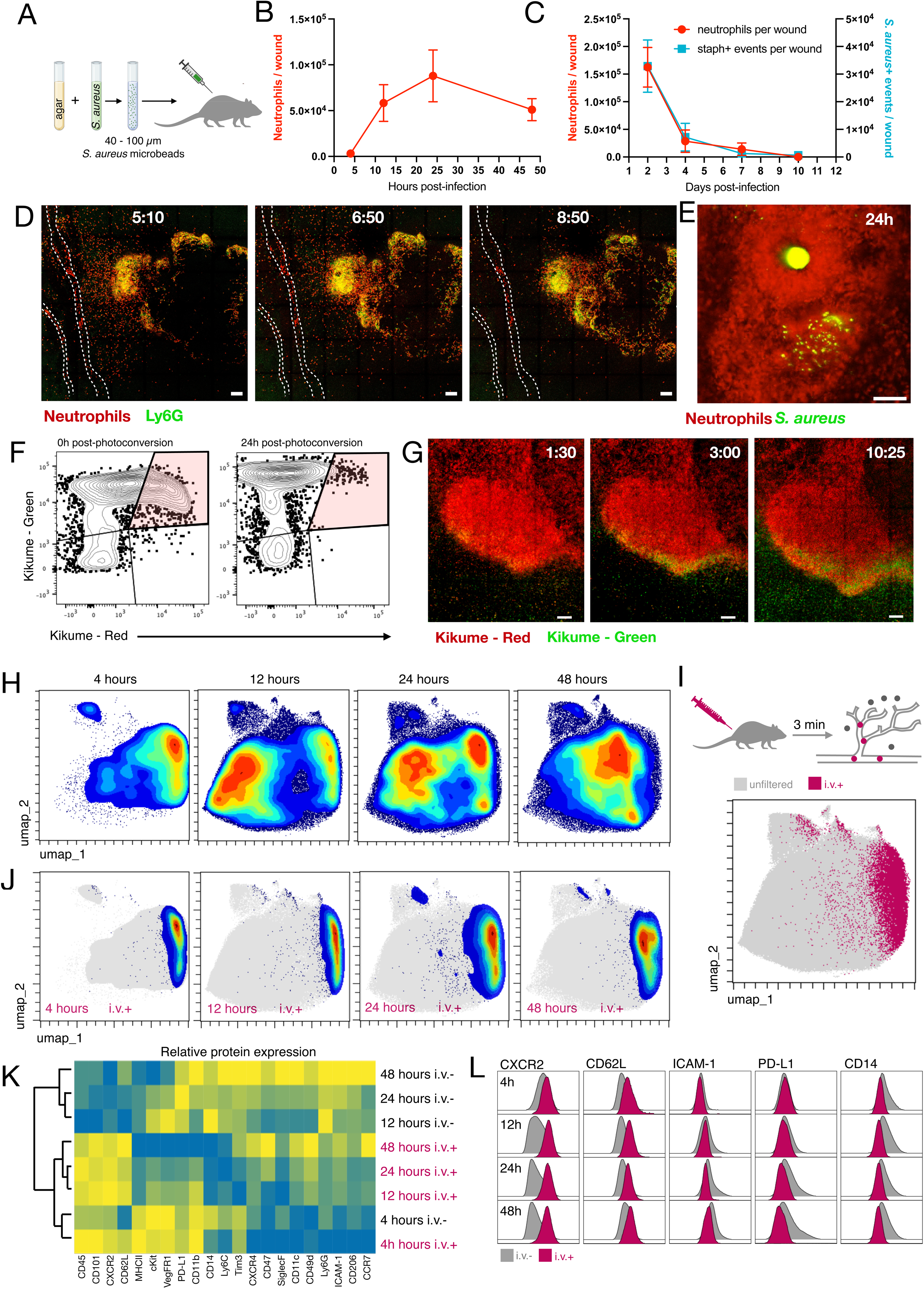
Neutrophil phenotypes are dynamic over the course of an infection response. A) Schematic representation of S. Aureus biofilm model of skin infection. B) Quantification by flow cytometry of neutrophil numbers per infectious wound over the initial 40 h of infection. N = 4-5 per time point. C) Quantification by flow cytometry of neutrophils and S. aureus+ events in infections over a 10-day time course. N = 5 mice per time point. D) Intravital microscopy of neutrophils (red, Catchup IVM-red) and Ly6G (green) in infectious wounds imaged between 5 and 9 hours post-infection. Scale bar = 100 μm E) Neutrophils (red, Catchup IVM-red) and S. aureus (Green, GFP) imaged in infectious wound at 24h of infection. Scale bar = 55 μm F) Flow cytometry data from Kikume mice representing Kikume-RED+ cells in the infection site 0h (left) and 24h (right) after photoactivation of the wound. G) Intravital microscopy of Kikume-RED (photoconverted) and KIkume-GREEN (non-photoconverted) cells within a 24h infection. Cells were photoactivated at t=0, and images taken at 1h30, 3h, and 10h25 after photoactivation. Scale bars = 100 μm H) UMAP projections of spectral flow cytometry data representing neutrophils from 4, 12, 24, and 48 hour skin infections. N = 4-5 mice per time point, concatenated I) Schematic representation of intravenous (i.v.) cell labeling to distinguish intravascular versus tissue-associated neutrophils in flow cytometry data. J) UMAP representation of intravenous+ cells (colored by density) and tissue-associated cells (gray) in 4h-48h infections. K) Hierarchically clustered heatmap of i.v.+ and i.v.- neutrophil phenotypic markers from 4h-48h infections. L) Histogram representations of flow cytometry data comparing selected marker expression in i.v.+ (pink) and tissue (i.v.-, gray) neutrophils from 4h-48h infections. Error bars = SEM.

To address the turnover of neutrophils within the site of infection, we used Kikume mice, which express a photoconvertible green fluorescent protein that becomes red when exposed to 405 nm wavelength (UV) light and maintains this signal for at least 24h [Fig. S1B](*20*). While we recovered a substantial population of Kikume-Red+ neutrophils from an infection photoconverted immediately before analysis, we did not detect appreciable levels of Kikume-Red+ neutrophils at 24h after photoconversion [Fig. 1F], indicating high levels of neutrophil turnover within the infection site. Indeed, cells with a neutrophil signature (CD11b-hi Ly6G-hi) were found to accumulate within the dead cell gate via flow cytometry over the time course of infection [Fig. S1C], suggesting that neutrophil death likely contributes to this rapid turnover. To assess neutrophil influx to the wound, we employed intravital imaging of Kikume mice. At 24h after infection, the wound site was exposed to UV light, then imaged over the course of 10.5 hours. Kikume-Green+ cells, which were not present in the wound immediately following photoconversion, were seen trafficking to and integrating within the swarm of Kikume-Red+ neutrophils. The majority of these newly arrived cells stayed at the margin of the neutrophil swarm and mixed with the Kikume-Red+ neutrophils in the border region [Fig. 1G]. Together, these results demonstrate that neutrophils populate an infection site early following challenge and undergo rapid dynamics of turnover and accumulation throughout the peak of infection.

### Neutrophils are capable of rapid phenotypic adaptation in an infection site

We next assessed how these dynamics corresponded with neutrophil phenotypes during the first days of an infection. To address this, we employed a 23-color spectral flow cytometry panel [Table S1] and analyzed neutrophil phenotypes at 4, 12, 24, and 48 hours after infection [Fig S1D-E]. Using unbiased dimensionality reduction, we observed that the neutrophil landscape shifted appreciably over this time course [Fig. 1H]. By including an intravascular label, we were able to distinguish neutrophils that were present within vessels (i.v. CD45+, or i.v.+) versus those that had extravasated into the wound site (i.v. CD45-, or i.v.-). This revealed that i.v.+ neutrophils occupied a similar area of UMAP across all time points analyzed, in contrast to the shifting phenotypic landscape seen within the tissue [Fig. 1I-J]. These data supported that neutrophil phenotypic diversity was largely a feature of extravascular neutrophils compared to those in the circulating pool.

Hierarchical clustering revealed that i.v.+ neutrophils were indeed more similar to each other, compared to phenotypes adopted in the i.v.- fractions across time points, with the exception of 4h i.v.- neutrophils, which presumably had insufficient time to change significantly from their i.v.+ counterparts [Fig. 1K]. Considering these indications of major differences between intravascular and extravasated neutrophils, we analyzed changes in several markers between the circulation and tissue over time [Fig. S1E]. We found that neutrophils both up- and down-regulated various proteins upon exiting the blood stream. Notably, we found substantial down-regulation of CXCR2 and CD62L, and up-regulation of ICAM-1, CD14, and PD-L1 [Fig. 1L]. Overall, these data indicate that, while neutrophils are relatively short-lived within the wound, they nonetheless undergo significant phenotypic specification in the context of an infection, and that these adaptations are markedly absent from the circulating pool of neutrophils.

### A novel population of CD101-lo mature neutrophils is associated with the infection response

One marker that changed dramatically over time during the infection was CD101, which was highest at 4h after infection, then appeared bimodal in expression at 12, 24, and 48h [Fig. 2A]. Comparing intravascular and extravasated neutrophils, we noted that i.v.+ neutrophils were almost entirely CD101-hi, while those in the tissue were both CD101-hi and CD101-lo [Fig. 2B], indicating potential loss of CD101 on extravasated cells. Since neutrophils are thought to up-regulate CD101 with maturation in a unidirectional manner(*21*), we sought to interrogate whether CD101-lo neutrophils in the tissue were were younger or less mature than those in the CD101-hi gate. Using EdU pulsing, which labels only proliferative neutrophil progenitors within the bone marrow (BM), we assessed relative ages of neutrophil subpopulations based on EdU signal accumulation [Fig. 2C]. In the bone marrow, we observed the expected kinetics of EdU labeling—EdU+ cells peaked in the BM CD101-lo gate 36-48h after pulsing, and subsequently in the BM CD101-hi gate 48-60h after pulsing. EdU signal then accumulated in the blood neutrophil gate (60+h after pulsing), reflecting the known maturation trajectory of neutrophils [Fig. 2D] (*21*). When we analyzed the emergence of EdU+ neutrophils in the skin, we found that the EdU signal accumulated in CD101-hi neutrophils with slightly more rapid dynamics than in CD101-lo neutrophils [Fig. 2D], in contrast to the dynamics the bone marrow. These data are consistent with CD101-hi neutrophils in the skin being slightly younger than the CD101-lo population, rather than CD101-lo neutrophils representing a younger population cells.

**Figure 2.**
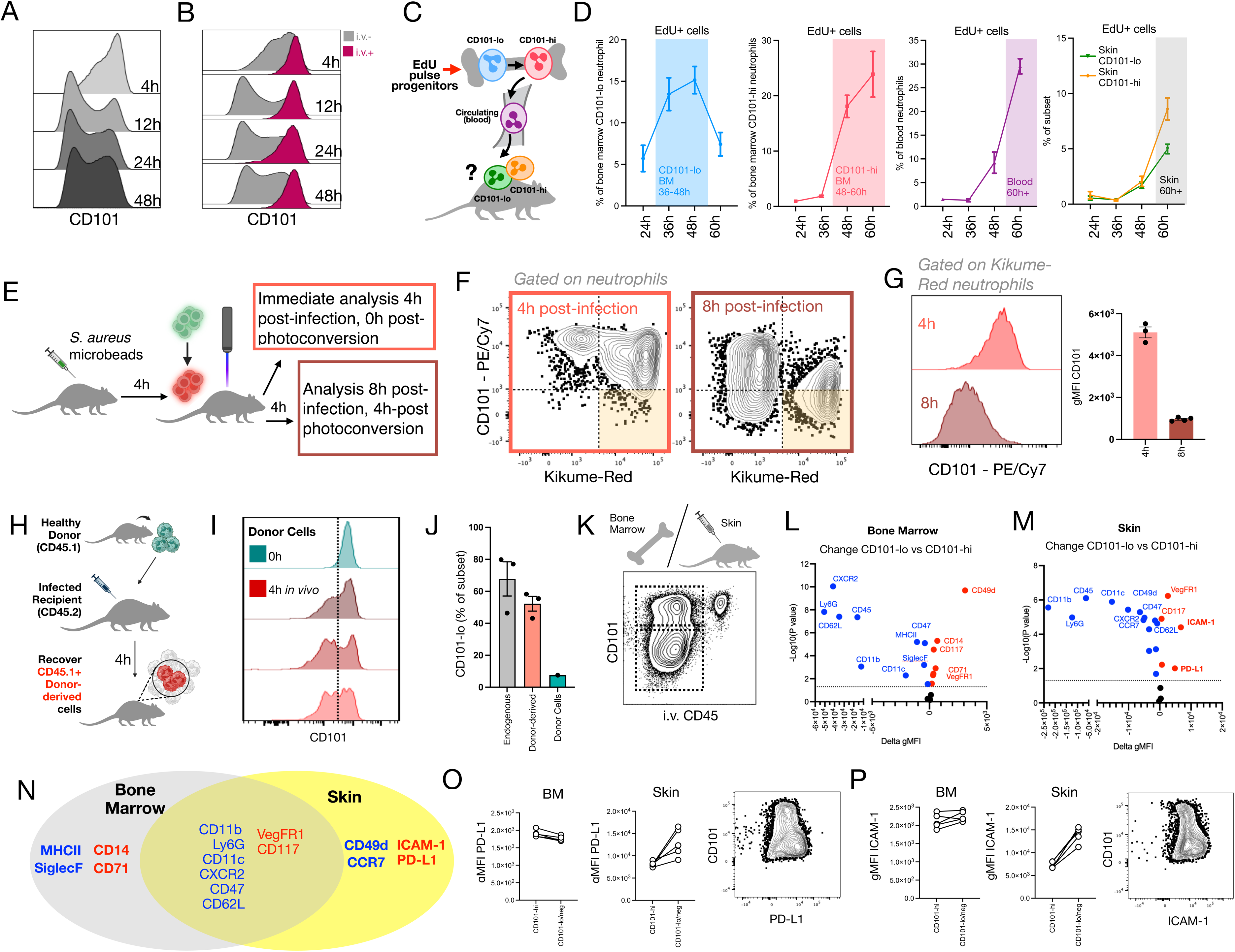
A novel population of CD101-low mature neutrophils is associated with infection response. A) Histogram representations of CD101 expression in neutrophils from 4h, 12h, 24h, and 48h infections. B) Histogram representations of CD101 expression in neutrophils from i.v.+ versus i.v.- compartments of h, 12h, 24h, and 48h infections. C) Schematic representation of EdU experimental design. D) Quantification of flow cytometry data depicting EdU positivity as a proportion of various neutrophil subsets. From left to right, CD101-lo bone marrow neutrophils (blue), CD101-hi bone marrow neutrophils (pink), blood neutrophils (purple), and CD101-hi (orange) or CD101-lo (green) skin infection-associated neutrophils. N = 4-5 mice per data point. E) Schematic depiction of adoptive transfer scheme, wherein blood neutrophils of CD45.1 congenic mice were isolated and transferred into infection sites of CD45.2 recipient mice, and analyzed 4h later for phenotypic traits. I) Histogram representations of flow cytometry data depicting CD101 expression in CD45.1 donor neutrophils (top, teal) and CD45.1 neutrophils recovered 4h after adoptive transfer (lower, red), each red histogram representing cells from an individual recipient mouse. J) Quantification of the proportion of endogenous CD45.2 (gray) and donor-derived CD45.1 (red) neutrophils falling into the CD101-lo gate. K) Schematic representation of analysis to compare CD101-hi to CD101-lo i.v.- neutrophils from either bone marrow or skin infections. L-M) Volcano plot representation of phenotypic markers and how they differ between CD101-lo and CD101-hi populations in bone marrow (L) and skin infections (M). Delta gMFI values represent averages over N = 5 mice. N) Venn-diagram representing markers that are decreased (blue) or increased (red) in expression within the CD101-lo gate, compared to the CD101-hi gate, from each tissue. O-P) Pair-wise comparisons of PD-L1 (O) or ICAM-1 (P) expression in CD101-hi versus CD101-lo subpopulations of neutrophils from bone marrow (left) and skin infections (middle), and representative flow cytometry data depicting PD-L1 expression in infection associated neutrophils (right). Error bars = SEM, statistics = t-test for P values shown in volcano plots.

We next directly interrogated whether CD101-hi mature neutrophils were losing CD101 expression after entering the tissue. To this end, we again used Kikume mice. Wounds were exposed to UV light at 4h after infection [Fig. 2E], a time point when all neutrophils in the tissue have a CD101-positive signature [Fig. 2A]. Analysis immediately after photoconversion showed that nearly all Kikume-Red+ neutrophils were indeed CD101+ [Fig. 2F]; however four hours later (8h post-infection), many of the Kikume-Red+ cells had shifted to the CD101- gate [Fig. 1F]. Gating on Kikume-Red+ neutrophils revealed discrepant expression of CD101 between the 4h and 8h infections [Fig. 2G]. These data indicate that CD101 is lost from neutrophils upon exiting the circulation and arriving in an infection site, corresponding to the period of swarm initiation captured by intravital microscopy between 5h-9h post-infection [Fig. 1D].

To address whether mature, circulating neutrophils from a naive mouse could phenotypically adapt in this manner upon exposure to an infection environment, we isolated blood neutrophils from CD45.1 donor mice and injected them directly into 24h infections of CD45.2 congenic recipient mice. After 4h, we recovered the transferred cells to assess CD101 expression [Fig. 2H]. We found that CD45.1+ neutrophils had indeed down-regulated CD101 compared to levels expressed before injection [Fig. 2I], with a similar proportion of CD45.1 (donor) and CD45.2 (endogenous) cells found in the CD101-lo gate, in contrast to the donor cells which were all CD101-hi before infection [Fig. 2J]. These data also supported that the act of extravasation was not necessary for CD101 down-regulation. Using unbiased dimensionality reduction with a simplified spectral flow cytometry panel [Table S1], we saw that the donor-derived CD45.1+ neutrophils recovered from the infection clustered among the endogenous cells rather than among naive donor cells [Fig. S2A], suggesting that the donor cells adapted their phenotype to align moreso with the endogenous cells across other markers as well. Together these findings indicate that mature, CD101-hi neutrophils rapidly lose CD101 expression upon exiting the circulation and entering a site of infection, and that CD101-lo neutrophils found within an infection are not younger than CD101-hi neutrophils in the same site.

We next assessed phenotypic traits of mature CD101-lo neutrophils from infection sites and immature CD101-lo neutrophils from the bone marrow. We compared CD101-lo cells from each tissue to CD101-hi counterparts, setting the CD101-hi cells as local, canonical mature neutrophil controls [Fig. 2K]. In the bone marrow, CD101-lo neutrophils were depleted for markers such as Ly6G, CD11b, CXCR2, CD62L, and CD11c, while CD117, CD49d, and CD71 were enriched, as expected [Fig. 2L]. In the skin, several of these patterns were reproduced, including lower expression of Ly6G, CD11b, CXCR2, CD62L, and CD11c, and increased expression of CD117 [Fig. 2M], suggesting some level of apparent phenotypic “de-differentiation” of the CD101-lo infection-associated state. However, some phenotypic changes distinguished CD101-lo skin neutrophils from those in the BM, including a reduced rather than increased expression of CD49d, and up-regulation of markers including PD-L1 and ICAM-1 [Fig. 2M-P]. These data support that CD101-lo converted mature neutrophils in the skin are phenotypically distinct from CD101-lo immature neutrophils in the bone marrow.

### CD101-lo PD-L1-hi ICAM-1-hi mature neutrophils represent a phagocytic “post-effector” neutrophil state

Artificially subsetting neutrophils into i.v.+ neutrophils, i.v.- CD101-hi neutrophils, and i.v.-CD101-lo neutrophils [Fig. 3A], we confirmed that the PD-L1-hi ICAM-1-hi state was enriched in the i.v.- CD101-lo gate [Fig. 3B], and observed that this population became more pronounced over time [Fig. 3C]. Using fluorescent *S. aureus*, we assessed pathogen binding, and found that CD101-lo neutrophils were enriched for *S. aureus* positivity [Fig. 3D], which was further enhanced in the PD-L1-hi ICAM-1-hi gate [Fig. 3E]. Using the UMAP projection of neutrophils from the two-day time course [Fig. 1H] to unbiasedly identify patterns in our dataset, we identified that side scatter (SSC-A), a metric for neutrophil granularity, correlated with the pattern of CD101 expression [Fig. 3F-G]. Neutrophils are granulocytes, and several of their key effector mechanisms involve releasing granular contents, which leads to a reduction in cellular granularity. This raised the possibility that CD101-lo neutrophils represented a degranulated, post-effector neutrophil state. We therefore tested whether key molecular pathways involved in degranulation would influence CD101-lo mature neutrophils. To this end, mice were treated with one of three inhibitors, targeting either NADPH oxidase, Rho kinase, or several proteases, or a saline control. All three interventions affected side scatter, indicating an impact on degranulation [Fig. 3H]. Although there was no significant affect on presence of the CD101-lo neutrophil state [Fig. S3A], emergence of the PD-L1-hi ICAM-1-hi CD101-lo state was blunted in each setting [Fig. 3I-J, Fig. S3B]. Together, these data support that specification of the CD101-lo neutrophil state in the infectious wound is tightly linked to key effector responses including pathogen binding and degranulation.

**Figure 3.**
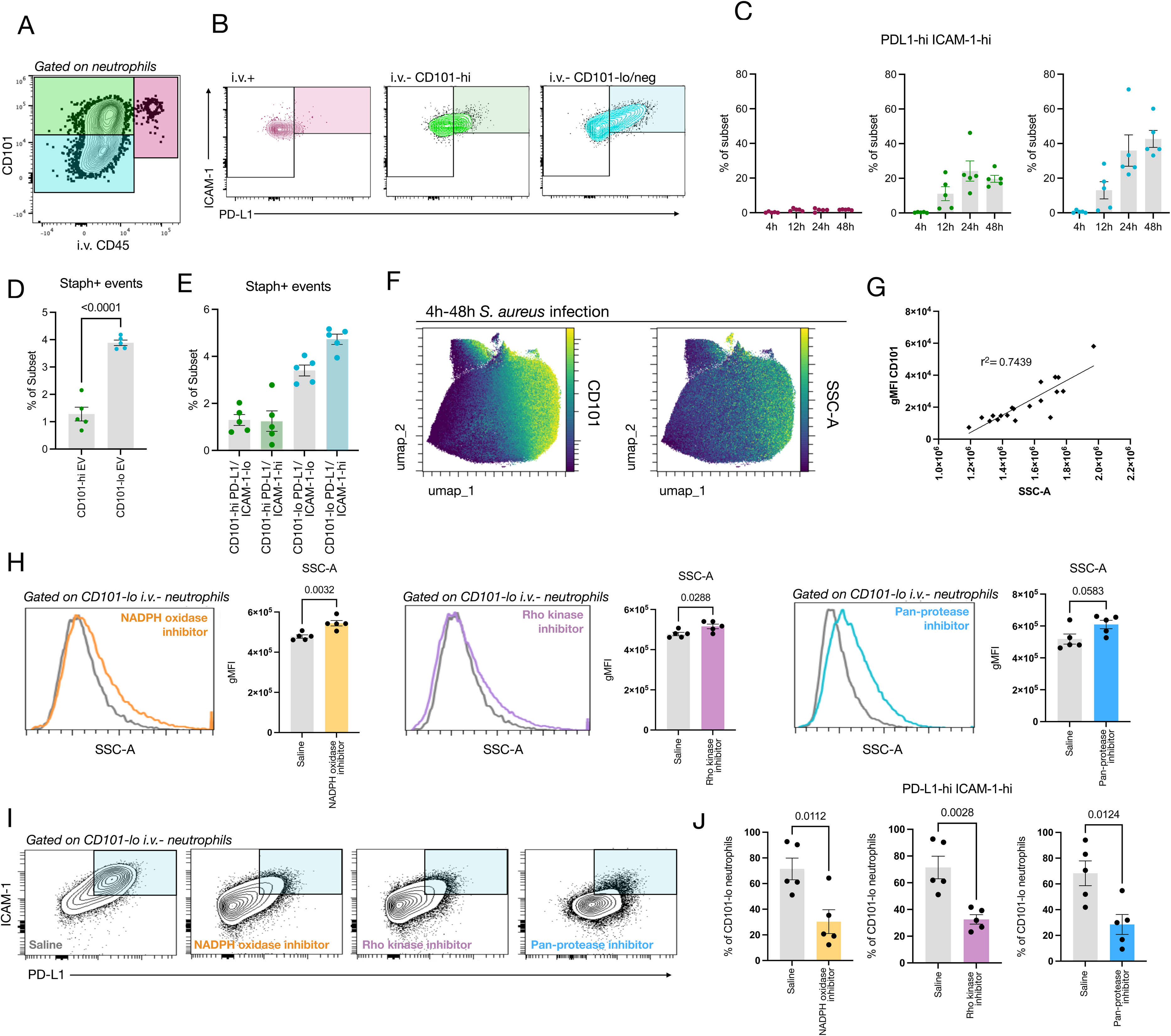
Differentiation of CD101-lo PD-L1-hi ICAM-1-hi mature neutrophils requires key effector processes. A) Schematic depicting artificial subsetting of neutrophils into i.v.+ (pink), i.v.-CD101-hi (green) and i.v.- CD101-lo (blue) subpopulations. B) Flow cytometry data depicting the distribution of each subpopulation, as in A, according to parameters of PD-L1 and ICAM-1 expression. C) Quantification of flow cytometry data as in B. N = 5 mice per group. D-E) Quantification of fluorescent S. Aureus signal within i.v.- CD101-hi and i.v.- CD101-lo subpopulations (D), and PD-L1-hi ICAM-1-hi versus PD-L1-lo ICAM-1-lo subgates (E), within neutrophil from 24h infections. N = 5 mice. F) UMAP representation of neutrophils from 4h-24h infections, colored by fluorescence intensity for CD101 (left) and side scatter (SSC-A, right) parameters. N = 4-5 mice per time point, concatenated. G) CD101 expression plotted against granularity (side scatter, SSC-A) for neutrophils from mice across the 4h-48h time course of infection; each point represents one mouse. H) Histogram representations and quantification of granularity (side scatter, SSC-A) in infection-associated neutrophils from mice treated with NADPH oxidase inhibitor, Rho kinase inhibitor, or Pan-protease inhibitor, or the relevant saline-treated control. Saline controls from NADPH oxidase and Rho Kinase inhibitors are equivalent as these experimental conditions were tested in tandem. N = 5 mice per condition. I) Flow cytometry data depicting the distribution of infection-associated neutrophils in the i.v.- CD101-lo gate per PD-L1 and ICAM-1 expression. J) Quantification of the proportion of CD101-lo i.v.- neutrophils within the PD-L1-hi ICAM-1-hi gate for each condition. N = 5 mice per group. Error bars = SEM, statistics = t-test, P values shown.

### Immature neutrophils can phenotypically mimic mature neutrophils, but mature neutrophils are required for optimal host defense

While the CD101-lo neutrophils populating skin infections were found to be mature cells, we wondered whether CD101-lo immature neutrophils would be similarly proficient in host defense. Indeed, it has recently been suggested that immature neutrophils may adapt their program similarly to mature neutrophils upon arrival to a tissue site in the setting of cancer(*22*), so we asked whether the same was true in the context of an infection. To this end, we took two approaches to enrich for immature neutrophils during the host defense response. First, we treated mice with G-CSF in order to enrich the circulating pool of neutrophils for immature neutrophils from the bone marrow [Fig. 4A]. Unbiased dimensionality reduction revealed that circulating neutrophils in G-CSF-treated mice were phenotypically different from untreated mice [Fig. 4B]. However, within the infection site, a similar distribution of neutrophils from each condition was observed [Fig. 4C]. Indeed, within the i.v.+ fraction of neutrophils, 14/21 phenotypic markers tested showed significantly different expression levels between treated and untreated mice, while i.v.- neutrophils showed only 4 markers discrepantly expressed [Fig. 4D], indicating phenotypic convergence of neutrophils within the tissue from each condition. While G-CSF-treated i.v.+ neutrophils showed much lower CD101 expression than those in control mice, reflecting mobilization of a CD101-lo immature neutrophil population from the bone marrow, within the wound CD101 expression displayed a remarkably similar pattern between conditions [Fig. 4E, S4A]. Functionally, however, neutrophils from G-CSF-treated mice had a limited capacity to control infection compared to untreated mice, despite similar wound size and neutrophil influx [Fig. 4E].

**Figure 4.**
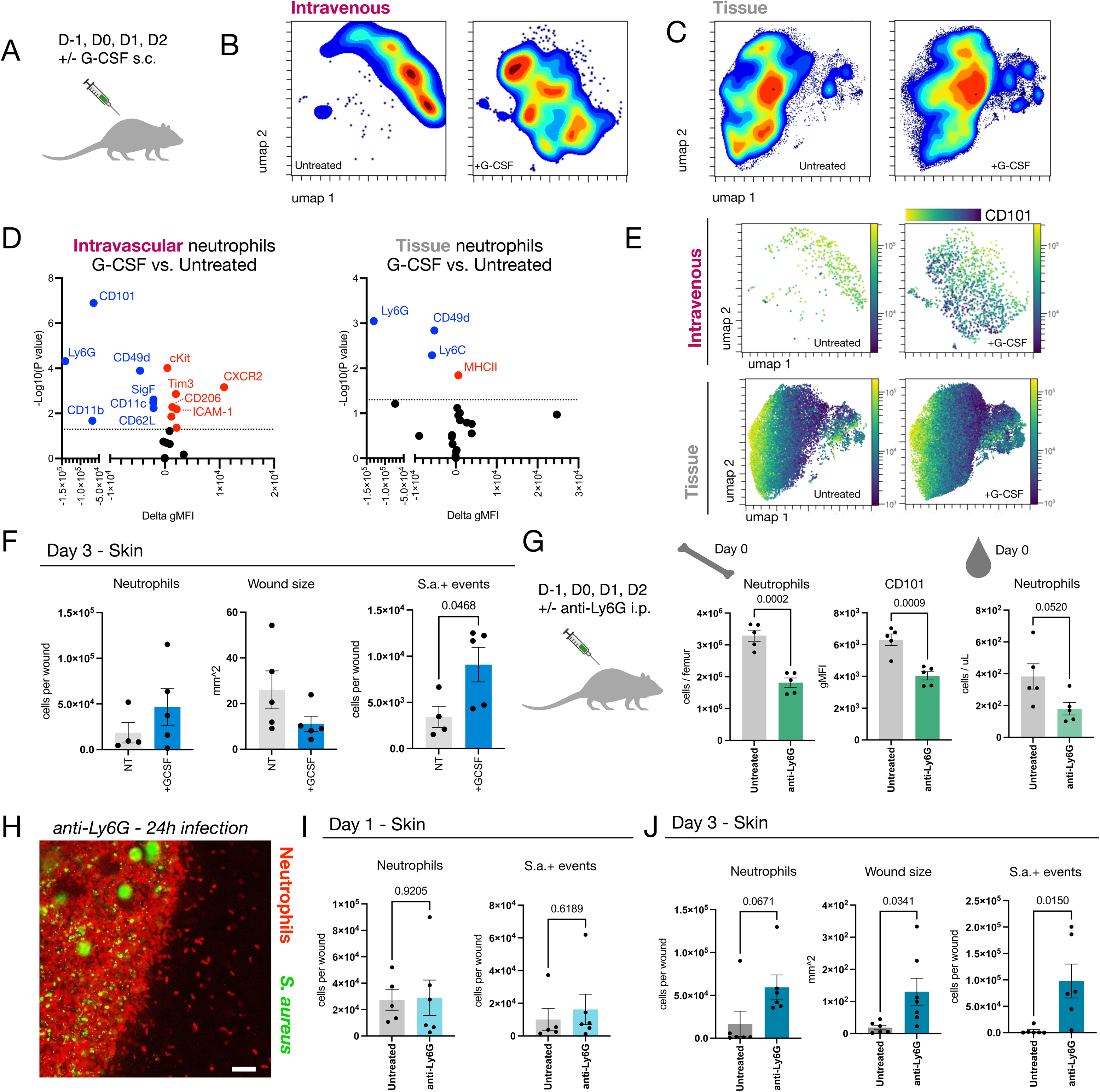
Immature neutrophils phenotypically mimic mature neutrophils but are deficient in host defense. A) Schematic depicting G-CSF treatment regimen for experimental mice. B-C) UMAP representation of spectral flow cytometry data depicting intravascular (B) or tissue-associated (C) neutrophils from 72h skin infections of control mice versus those treated subcutaneously with G-CSF. N = 4-5 mice per group. D) Volcano plot representation of differences neutrophil phenotypic markers between control and G-CSF-treated mice, from intravascular (left) and tissue-associated (right) neutrophil compartments. Data averaged from N = 4-5 mice. E) UMAP representations of spectral flow cytometry data from intravenous (top) and tissue-associated (bottom) neutrophils in control (left) versus G-CSF-treated (right); colored by fluorescence intensity for CD101. F) Quantification of flow cytometry data depicting neutrophil numbers within infection sites (left), infectious wound size (middle), and *S. aureus*+ events per wound (right) from infections of control versus G-CSF-treated mice. For B-F, one outlier in control group was excluded on basis of ROUT outlier identification test (Q = 1%). G) Schematic representation (left) and flow cytometry data quantification (right) for mice treated with i.p. anti-Ly6G antibody versus control mice. Quantification of bone marrow neutrophil numbers (middle left) and their CD101 expression (middle right), as well as blood neutrophil numbers (right) are presented. N = 5 mice per group. H) Intravital microscopy depicting neutrophils (red, Catchup IVM-red) and *S. aureus* (green) in an infection site from a mouse treated with anti-Ly6G antibody. Scale bar = 50 μm I) Quantification of flow cytometry data depicting neutrophil and *S. aureus* numbers in 24h infections from mice treated with anti-Ly6G or control mice on days. N = 5-6 mice per group. J) Quantification of flow cytometry data depicting neutrophil and *S. aureus* numbers in 72h infections (left, right) and wound sizes (middle) from mice treated with anti-Ly6G or control mice. N = 6 mice per group. Error bars = SEM, statistics = t-test for bar charts and volcano plots, P values shown.

Second, we asked whether simply depleting the most mature neutrophils would yield a similar outcome. We thus treated mice with anti-Ly6G at a dose of 200 ug i.p. It has recently been suggested that treatment with anti-Ly6G poorly depletes neutrophils, enriching for less mature neutrophils(*23, 24*), which we confirmed [Fig. 4G]. While neutrophil numbers were altered in the blood of anti-Ly6G-treated infected mice at 24h [Fig. S4B], we found no differences in wound homing, infection control, or overall inflammation at this time point [Fig. 4H-I, Fig. S4B]. However, by 3d post-infection, anti-Ly6G-treated mice failed to control their infections, demonstrating highly inflamed, neutrophil-infiltrated lesions and an elevated burden of *S. aureus* [Fig. 4J]. Together, these data support that although immature neutrophils are capable of phenotypic specification, they are intrinsically different from mature neutrophils and less capable of host defense.

### Neutrophil phenotypic adaptation wanes during the resolution phase of the inflammatory response

We next investigated the trajectory of neutrophil phenotypic specification during the resolution phase of the infection-associated inflammatory response. Using our spectral flow cytometry panel to compare skin neutrophils during peak (days 1, 2) and resolution (days 3, 4) phases, we found significant differences between each time point assessed [Fig. 5A]. Similar to the early time course (4h-48h) [Fig. 1H-J], we again observed that i.v.+ neutrophils changed minimally across the resolution phase [Fig. 5B]. The CD101-lo neutrophil population, as a proportion of i.v.- neutrophils, diminished between days 1-4 [Fig. 5C], and this corresponded with progressively blunted reductions in granularity (SSC-A) in the i.v.- neutrophil populations [Fig. 5D]. Adaptation of neutrophil phenotypes from the i.v.+ to i.v.- fractions were also progressively blunted, with more limited phenotypic changes evident at days 3 and 4 post-infection [Fig. 5E]. This was confirmed with a hierarchically clustered heatmap, showing that day 1 and 2 i.v.- neutrophils were distinct from i.v.+ subsets, but that the i.v.- populations on days 3 and 4 were highly similar to i.v.+ cells [Fig. 5F]. Finally, as the levels of neutrophils in the wound diminished, so too did prevalence of the CD101-lo PD-L1-hi ICAM-1-hi post-effector state [Fig. 5G]. These data indicate that neutrophil specification within the host defense niche becomes restrained as inflammation resolves.

**Figure 5.**
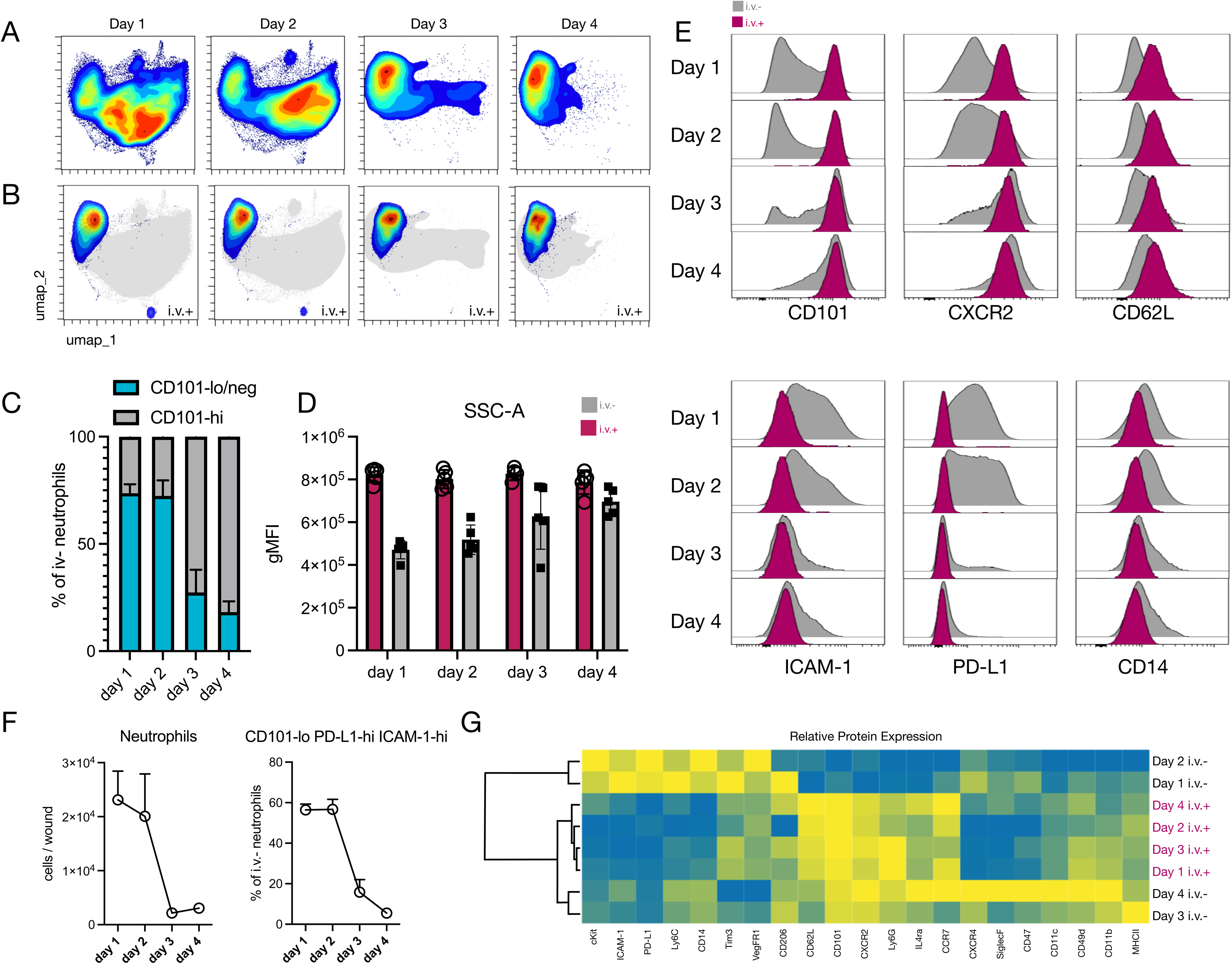
Neutrophil phenotypic specification wanes during inflammation resolution. A) UMAP projections of spectral flow cytometry data representing neutrophils from 1, 2, 3, and 4 day skin infections. N = 4-5 mice per time point, concatenated B) UMAP representation of intravenous+ cells (colored by density) and tissue-associated cells (gray) in 1-4 day infections. N = 4-5 mice per time point, concatenated. C) Quantification of flow cytometry data depicting the proportion of CD101-lo/ negative versus CD101-hi neutrophils in the i.v.- compartment of skin infections over a 1-4 day time course. N = 4-5 mice per time point. D) Quantification of flow cytometry data depicting neutrophil granularity (side scatter, SSC-A) in i.v.+ and i.v.- compartments of day 1-4 skin infections. N = 4-5 mice per group. E) Histogram representations of flow cytometry data comparing selected marker expression in i.v.+ (pink) and tissue (i.v.-, gray) neutrophils from 1-4 day infections. F) Quantification of flow cytometry data depicting neutrophil numbers (left) and the proportion of CD101-lo PD-L1-hi ICAM-1-hi neutrophils in the i.v.- compartment (right) of day 1-4 skin infections. N = 4-5 mice per time point. G) Hierarchically clustered heatmap of i.v.+ and i.v.- neutrophil phenotypic markers from 1-4 day infections.

### Neutrophil phenotypic adaptation is challenge- and not source-dependent

We next investigated how different acute barrier challenges in the skin might impact neutrophil phenotypes. To this end, we compared the setting of a *S. aureus* biofilm infection to an aseptic wound induced by making a small excision of skin. These two challenges induced comparable levels of neutrophil infiltration at 24h after challenge [Fig. 6A]. Using spectral flow cytometry and unbiased dimensionality reduction, plus artificial clustering to regionally annotate the UMAP [Fig. S5A], we found that neutrophils from each setting presented in a unique pattern across the neutrophil landscape [Fig. 6B, Fig. S5A-B]. Cluster representation differed significantly between conditions [Fig. 6C-D], but we again observed minimal differences in the distribution of i.v.+ neutrophils across conditions, in contrast to the i.v.- neutrophils in the tissue [Fig. 6E, Fig. S5C]. Within the wound, expression or markers including CD101, CXCR2, CD62L, and PD-L1 were notably discrepant between settings [Fig. 6F]. Considering the major differences in CD101 and CXCR2 expression between infection and aseptic conditions, we asked whether a single mouse experiencing both challenges on contralateral flanks would still show divergent neutrophil phenotypes in each site. Indeed, in this setting, infections harbored neutrophils with low expression of CD101 and CXCR2, while aseptic wounds in the same mouse retained higher CD101 and CXCR2 expression [Fig. 6G-H], supporting that phenotypic changes from blood to tissue are independent of the circulating neutrophil pool and rather dependent on the local tissue environment. Overall, these data support that neutrophil phenotypic specification is challenge-dependent and not dependent on different source pools of neutrophils being available in the circulation.

**Figure 6.**
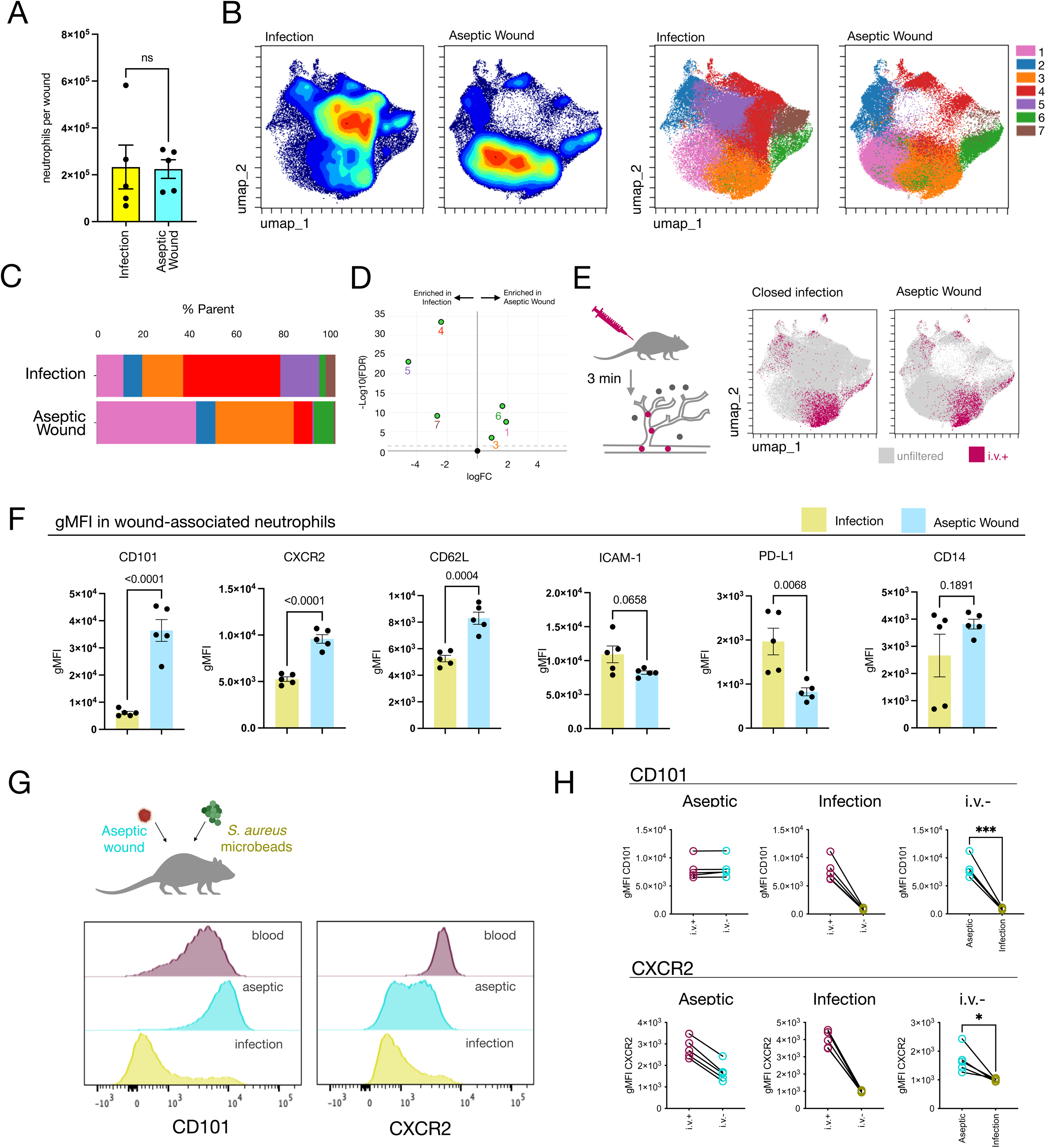
Neutrophil phenotypic adaptation is challenge- and not source-dependent. A) Quantification of flow cytometry data depicting numbers of neutrophils in skin infections compared to aseptic excisional wounds. N = 5 mice per condition. B) UMAP projections of spectral flow cytometry data representing neutrophils from 24h infections versus aseptic wounds (left), and clustering representation of spatial regions or the UMAP (right). N = 5 mice per condition. C) Bar plots representing the relative proportion of each cluster (1–7) within the total neutrophils from each condition. D) edgeR analysis of differential cluster representation between infection and aseptic wound conditions. E) Schematic depiction (left) and UMAP projections (right) of intravenous (i.v.+) neutrophils (pink) overlaid on all infection-associated neutrophils (gray) for infection (left UMAP) and aseptic wound (right UMAP) conditions. F) Quantification of flow cytometry data representing fluorescence intensity among selected neutrophil markers in infections versus aseptic wounds. G) Experimental schematic (top) and histogram representations of flow cytometry data (bottom) from contralateral 24h infections versus aseptic wound challenges. H) Pairwise comparisons of CD101 and CXCR2 expression from contralateral 24h infections and aseptic wounds.

### Neutrophil responses differ between two challenges with the same pathogen

Considering the striking differences in neutrophil specification between biofilm infection and aseptic wounding, we next asked if neutrophils would respond similarly or differently to biofilm versus planktonic (free-floating) *S. aureus* challenge [Fig. 7A]. Since *S. aureus* in biofilms are resistant to phagocytosis, and in our model associated with foreign body structures, we anticipated that this comparison could reveal neutrophil traits associated with a frustrated neutrophil host defense response. Initially, we found that planktonic *S. aureus* challenge required a 20-30-fold higher inoculation dose of *S. aureus* to yield an appreciable neutrophil response in the skin [Fig. 7B]. Choosing the planktonic dose with neutrophil levels most similar to the biofilm model, we found that biofilm infections recruited more neutrophils per *S. aureus* (higher neutrophil / *S. aureus* ratio at 24h) [Fig. 7C]. Yet similar numbers of neutrophils were found to bind *S. aureus* in both settings [Fig. 7D], demonstrating that recruitment outscales the need for pathogen binding moreso in biofilm infection than with planktonic challenge. While neutrophils in the i.v.+ fraction had similar granularity (SSC-A) between settings, those encountering biofilm infection showed a much more striking reduction in granularity in tissue (i.v.-) [Fig. 7E], indicating a boosted degranulation response.

**Figure 7.**
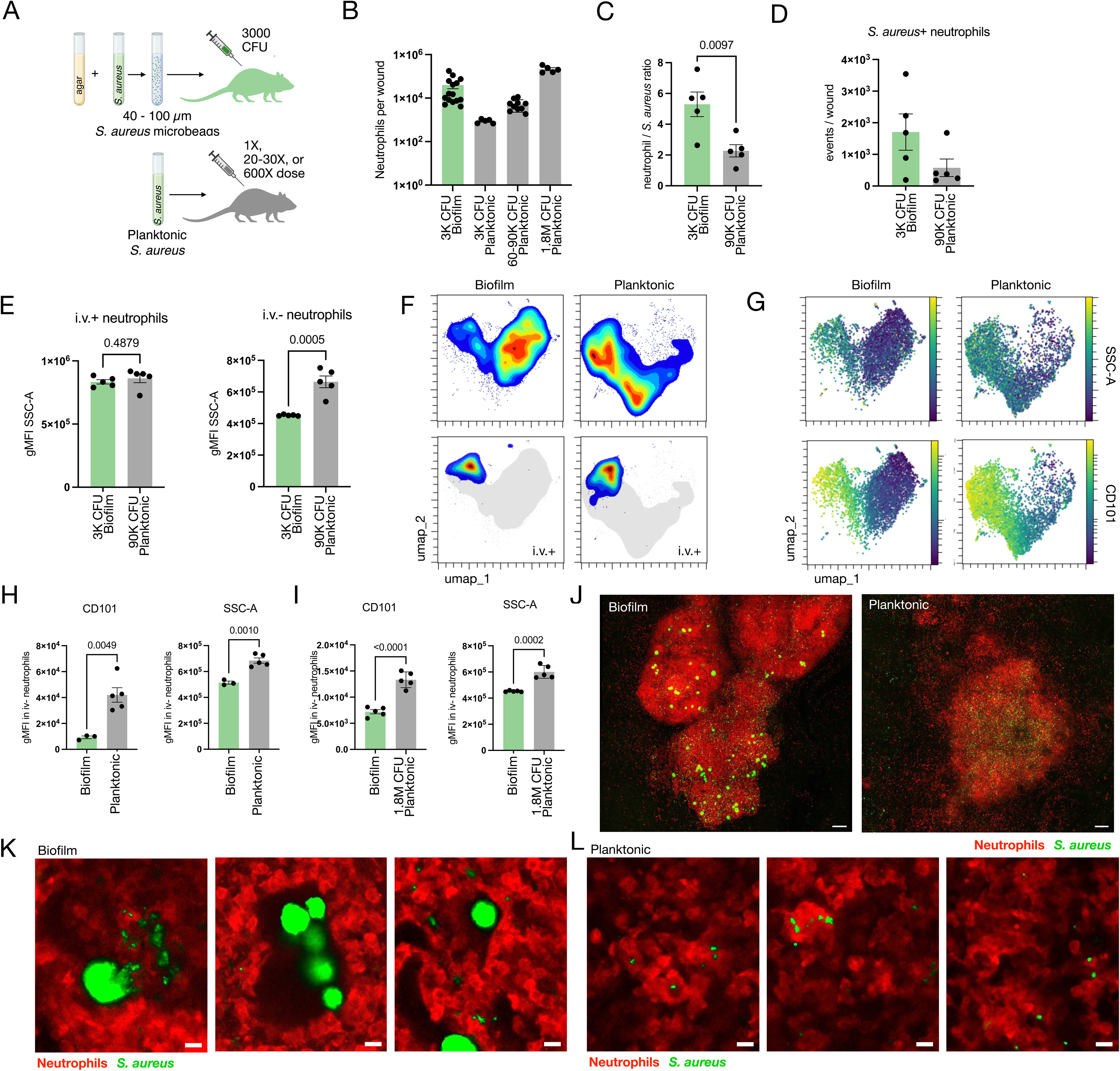
Neutrophil responses differ between biofilm- and planktonic-*S. aureus* challenge. A) Schematic depicting *S. aureus* biofilm (top) and planktonic (bottom) infection models. B) Quantification of flow cytometry data depicting neutrophil numbers across various infection MOIs and models. Data pooled from 3 experiments. C) Quantification of flow cytometry data depicting ratios of neutrophils to *S. aureus* within infections. D) Quantification of flow cytometry data depicting numbers of *S. aureus*+ events in biofilm and planktonic infections at the indicated MOIs. E) Quantification of fluorescence intensity reflecting neutrophil granularity (side scatter, SSC-A) in the intravascular (i.v.+, left) and extravascular (i.v.-, right) compartments of 24h biofilm versus planktonic infections. N = 5 mice per condition. F) UMAP representations of spectral flow cytometry data from mice with either biofilm (left) or 60K CFU MOI planktonic (right) infections, reflecting total tissue neutrophils (top colored, bottom gray) or the i.v.+ compartment (bottom colored) at 24h. N = 3-5 mice per condition. G) UMAPs as in F, colored to reflect fluorescence intensity for side scatter (SSC-A, top) or CD101 expression (bottom). N = 3-5 mice per condition. H-I) Quantification of flow cytometry data depicting CD101 (left) and side scatter (right) fluorescence intensity from mice as in F-G (H), or a separate cohort of biofilm versus planktonic infection-bearing mice (I). J) Intravital microscopy depicting neutrophils (red, Catchup IVM-red) and S. aureus (green) in day 1 biofilm versus planktonic infections. Scale bar = 100 μm. K-L) Intravital microscopy depicting neutrophils (red, Catchup IVM-red) and S. aureus (green) in day 1 biofilm (K) versus planktonic (L) infections. Scale bars = 10 μm. Error bars = SEM, statistics = t-test for bar charts and volcano plots, P values shown.

To compare the phenotypic profile of neutrophils between settings, we selected only mice with similar levels of neutrophils per wound from one experiment [Fig. S6A] and performed unbiased dimensionality reduction using our spectral flow cytometry panel. This revealed major differences in the neutrophil phenotypic landscape between settings, again largely restricted to the i.v.- fraction [Fig. 7F]. A striking difference was a depletion of the CD101-lo SSC-A-lo population in planktonic challenge [Fig. 7G-H]. A similar pattern was observed in a second experiment wherein wounds were not selected for by neutrophil numbers [Fig. S6B]. To address whether these findings could be due to the magnitude of the infection or inflammatory response, we compared the same parameters (CD101, SSC-A) in a high-dose (1.8 x 10^6^ CFU) planktonic challenge, which recruited more neutrophils than the biofilm model and harbored a higher pathogen burden at the 24h time point [Fig. S6C]. In this setting, CD101 expression and granularity were still higher in the planktonic compared to the biofilm response [Fig. 7I, Fig. S6D]. Intravital microscopy revealed that neutrophils generated swarms in both biofilm and planktonic infections [Fig. 7J]. However, while neutrophils in biofilm infections could access *S. aureus* that had been liberated from biofilms [Fig. 1E, 7K], they frequently failed to bind biofilm-associated pathogen, forming dense swarms around foreign body biofilm structures [Fig. 1E, 7K]. In contrast, *S. aureus* particles in planktonic infections were readily accessible to neutrophil binding [Fig. 7L]. These data support that a large or effector-response-resistant challenge recruits an enhanced neutrophil response, of which a highly degranulated, CD101-lo mature neutrophil state is a major feature.

## Discussion

In this study, we sought to analyze the intricacies of neutrophil diversity in an evolutionarily powerful setting: infection and acute injury to a barrier site. We chose a biofilm model of *S. aureus* skin infection, as this mimics the real-world setting of a low MOI foreign body-associated infection, recruits a robust neutrophil response, and resists clearance for up to 7+ days. In this context, we investigated dynamics and mechanisms underlying neutrophil phenotypic diversity, in order to establish a foundational picture of neutrophils’ native propensities for adaptation and specification within their environment. Ultimately, we aligned our findings with real-time readouts of neutrophil behaviors observed through intravital microscopy, together painting an integrated picture of the neutrophil behavioral dynamics and phenotypic landscape during host defense.

Our study supports that mature neutrophils are highly plastic when responding to an infectious challenge in the skin, altering several phenotypic markers after extravasation and during the swarming response. This phenotypic adaptation was underscored by significant levels of neutrophil turnover, was most striking during the peak of neutrophilic inflammation, and waned during inflammation resolution. We identified an unexpected population of CD101-lo mature neutrophils as early as 8h after infection initiation and observable throughout the peak of inflammation. CD101 up-regulation is canonically considered to progress unidirectionally with neutrophil maturation. While it has been suggested that mature neutrophils may down-regulate expression of CD101 *in vitro*(*25*), the definitive emergence of CD101-lo mature neutrophils *in vivo* has, to our knowledge, not been observed to date. Here, we conclusively show CD101 loss from previously CD101-hi cells, and confirm that CD101-lo neutrophils in the infection response are not younger than CD101-hi cells in the same tissue. These findings beg caution of those using CD101 as a marker for neutrophil maturity in the periphery, and position CD101 modulation as a key feature of some peripheral inflammatory responses.

Although CD101-lo mature neutrophils showed some evidence of phenotypic “de-differentiation” (CD117 up-regulation, Ly6G, CD11b, CXCR2 down-regulation), these cells were distinct from immature CD101-lo neutrophils in the bone marrow in that they up-regulated PD-L1 and ICAM-1 surface expression. They also demonstrated a dramatic reduction in side scatter, suggesting a degranulated state. Notably, the CD101-lo PD-L1-hi ICAM-1-hi subpopulation was dependent on neutrophil effector pathways and associated with pathogen binding. We thus position CD101-lo PD-L1-hi ICAM-1-hi neutrophils as a “post-effector” subpopulation within the wound site that remains capable of pathogen binding.

Neutrophil phenotypic specification in the skin was also observed in immature neutrophils, although these cells were deficient in host defense capacity compared to mature neutrophils. These data support the notion, raised recently in the context of cancer(*22*), that immature neutrophils are plastic and can adapt to their environment similarly to mature cells. However, their intrinsic abilities to defend against pathogen challenge were diminished.

We found that neutrophil adaptation was highly dependent on the nature of a challenge. An aseptic wound to the skin induced a similar magnitude of neutrophil infiltration, but neutrophil phenotypes differed dramatically in this setting—which we showed was caused by a difference in local environment, rather than an altered supply of neutrophils in circulation. Finally, we compared biofilm challenge to a planktonic *S. aureus* infection in the skin, which poses fewer physiological challenges to the neutrophil effector response (i.e. no foreign body structures, reduced pathogen hardiness). Planktonic pathogen elicited a severely dampened neutrophil response relative to MOI, requiring 20-30x more CFUs to recruit an appreciable neutrophil response in the skin. During this response there was sparse representation of the CD101-lo SSC-A-lo neutrophil state. As such, we posit that the CD101-lo, low granularity neutrophil profile reflects a frustrated neutrophil response to large or recalcitrant barrier challenge.

In conclusion, our results reveal dynamics and mechanistic nuances underlying observations of neutrophil diversity *in vivo*. By studying neutrophils in their evolved niche, we have shed light on the fundamental capacities of neutrophils to phenotypically adapt to their environment, holding significant implications for understanding neutrophil diversity across disease settings. In light of our findings, we predict that features of the neutrophil response such as tissue-associated half-life and cooperative behaviors like swarming should broadly have major bearing on neutrophil phenotype. Finally, our discovery of a novel, mature CD101-lo neutrophil population, associated with the response to large, recalcitrant challenge, highlights how specific neutrophil behaviors (e.g. dense swarming, degranulation) may map to discrete phenotypes *in vivo*.

## Methods

### Biofilm Infection Model

*S. aureus* was cultured overnight in brain heart infusion buffer then subcultured for 1h 45min, after which bacteria were washed in sterile HBSS without calcium and magnesium (HBSS-). Bacteria were diluted to an OD 660 of 0.17-0.18, mixed at a 1:10 v/v ratio with warm 1.5% BHI agar (in PBS), and added dropwise into an ice cold Tween 20 / mineral oil (1:100 v/v) bath and stirred with a magnetic stir bar for 30 minutes. Bacteria/agarose microbeads were then washed in sterile HBSS-10 times to remove residual mineral oil, passed through a 100 μm cell strainer, then isolated with a 40 μm cell strainer and washed a final time in sterile HBSS-. 3 separate 30 uL aliquots of microbeads were then dissociated in 300 uL of HBSS- by passage through a 30-gauge syringe and plated in dilution for CFU enumeration. Based on CFUs, microbeads were diluted to an injection dose of 3000 CFU / 30 uL, injected into the dorsal skin of the mouse using a 23-gauge needle.

### Aseptic wounding model

Mice were anesthetized under isoflorane, dorsal fur shaved, and sanitized using 70% ethanol. An approximately 5 mm diameter incision was made in the skin of the mouse using surgical scissors.

### Planktonic

*S. aureus* was cultured overnight in brain heart infusion buffer then subcultured for 1h 45min, after which bacteria were washed in sterile HBSS without calcium and magnesium (HBSS-). Bacteria were diluted to an OD 660 of 1.0. For 3K CFU MOI, bacteria were further diluted in sterile HBSS- to achieve the indicated MOI per 30 uL intradermal injection, which was verified via CFU enumeration

### Mice

Wild type and transgenic mice on the C57Bl/6 background were bred in-house or purchased from Jackson Labs. Mice aged ∼6.5-14 weeks were used for experiments. Transgenic / congenic strains: Kikume, Catchup iVM-red, CD45.1 (B6.SJL-*Ptprc^a^ Pepc^b^*/BoyJ, JAX cat#: 002014

### Intravital microscopy and image processing

Mice were anesthetized via intraperitoneal injection of ketamine/xylazine (200 mg/kg and 10 mg/kg respectively). A midline incision was made on the back skin and sutures attached in order to secure a flap of skin upon a cover slip for imaging. Mice were imaged using either a Leica upright SP8 MP, Leica inverted SP8 MP DIVE, or Leica inverted Stellaris MP DIVE system and either confocal or multiphoton imaging settings. Images were processed using Fiji (ImageJ) software.

### Tissue processing and flow cytometry

Mice were injected with 3 ug of fluorescent anti-CD45 antibody and 3 minutes later euthanized for tissue isolation. Skin samples containing infection or wound site were minced in a 2 mL volume of digestion buffer containing 5 mM EDTA, 3% FBS, and 0.8 mg / mL collagenase II in HBSS-. Samples were then rocked at 37-C for 75 minutes and passed through a 70 um cell strainer to generate a single cell suspension. Isolated cells were washed, stained with GhostRed 710 viability dye (Tonbo), blocked with Fc Block (BioXCell) and stained with fluorescent antibodies. Samples were acquired on either an LSRII (BD) or Cytek Aurora spectral flow cytometer.

For bone marrow, femurs were harvested and heads removed with a razor blade, then bone marrow was into FACS buffer using quick pulses of centrifugation. Harvested cells were subject to ACK lysis for up to 5 minutes and resuspended in FACS buffer for staining.

### Kikume mouse model

Kikume transgenic mice were infected with *S. aureus* as noted above. At the indicated time points, dorsal skin in the area of the infection was exposed to a 405 nm wavelength light via a handheld laser (Laserglow Technologies) for up to 5 minutes. For microscopy, Kikume bone marrow was used to reconstitute the hematopoietic compartments of irradiated C57BL/6 mice. Eight weeks later, infection was initiated as above, and 24h later the infection site was exposed and prepped for imaging, then directly exposed to 405 nm wavelength light via handheld laser and directly imaged. For figure S1B, bone marrow chimeric Kikume mice were given a focal thermal injury to the liver as previously described [Wang et al., Science, 2022]. 24h later the injury was exposed and photoactivated via handheld 405 nm wavelength laser, then imaged directly or 24h later using the Leica MP DIVE inverted system.

### EdU

Mice were injected with 1 mg EdU in 100 uL volume saline i.p. at either 60, 48, 36, or 24 hours before analysis. 24h before analysis, skin infections were initiated. EdU staining was performed with the Click-iT Plus EdU Alexa Fluor 647 Flow Cytometry Assay Kit from Invitrogen.

### Adoptive Transfer

CD45.1 donor mice were anesthetized under isoflorane and blood was isolated via cardiac puncture into a tube containing 50 mM EDTA. Blood was washed with 15 mM EDTA 1% BSA in HBSS and layered onto a 78%, 69%, 52% Percoll gradient then centrifuged at 1500 x g for 30 minutes at room temperature with brake set to level 2. Neutrophils at the 78% / 69% interface were collected, washed with HBSS/EDTA/BSA, subject to ACK lysis, then washed and resuspended in HBSS for enumeration. 1 million cells per mouse were injected directly into 24h skin infections of CD45.2 recipient mice, and 4 hours later infections were isolated for flow cytometry analysis.

### Molecular Inhibitors

Rho kinase (50 ug in 20 uL saline) and NADPH oxidase inhibitor (10 ug in 20 uL DMSO+saline) were injected directly to the infection site at 3h and 8h after infection initiation. Pan-protease inhibitor was injected directly to the infection site at (50 ug in 20 uL) approximately 4h and 8h after infection initiation. Infections were harvested and analyzed 1 day later in comparison to saline injected controls.

### G-CSF and anti-Ly6G treatment

For neutrophil targeting, mice were treated with 200 ug i.p. of anti-Ly6G (Clone 1A8, BioXCell) on days -1, 0, 1, and 2 of infections. For G-CSF treatment, mice were injected subcutaneously with 125 ug / kg of G-CSF on days -1, 0, and 1 and ∼90 ug / kg on day 2 of infections.

### Data analysis

Flow cytometry and spectral flow cytometry were analyzed using FlowJo (BD) and Omiq.ai. Spectral flow cytometry data was collected on a Cytek Aurora (5-laser) nad spillover/compensation was adjusted in FlowJo before export to omiq.ai for further analysis. UMAP projections in omiq.ai were based on the majority of fluorescent parameters excluding intravenous antibody and fluorescent reporter *S. aureus* (See supplemental table). For FlowSOM clustering, Comma-separated k values were set to 30 for overclustering, and clusters were merged manually to unify spatial regions of the UMAP.

## Figure Legends

**Figure S1.**
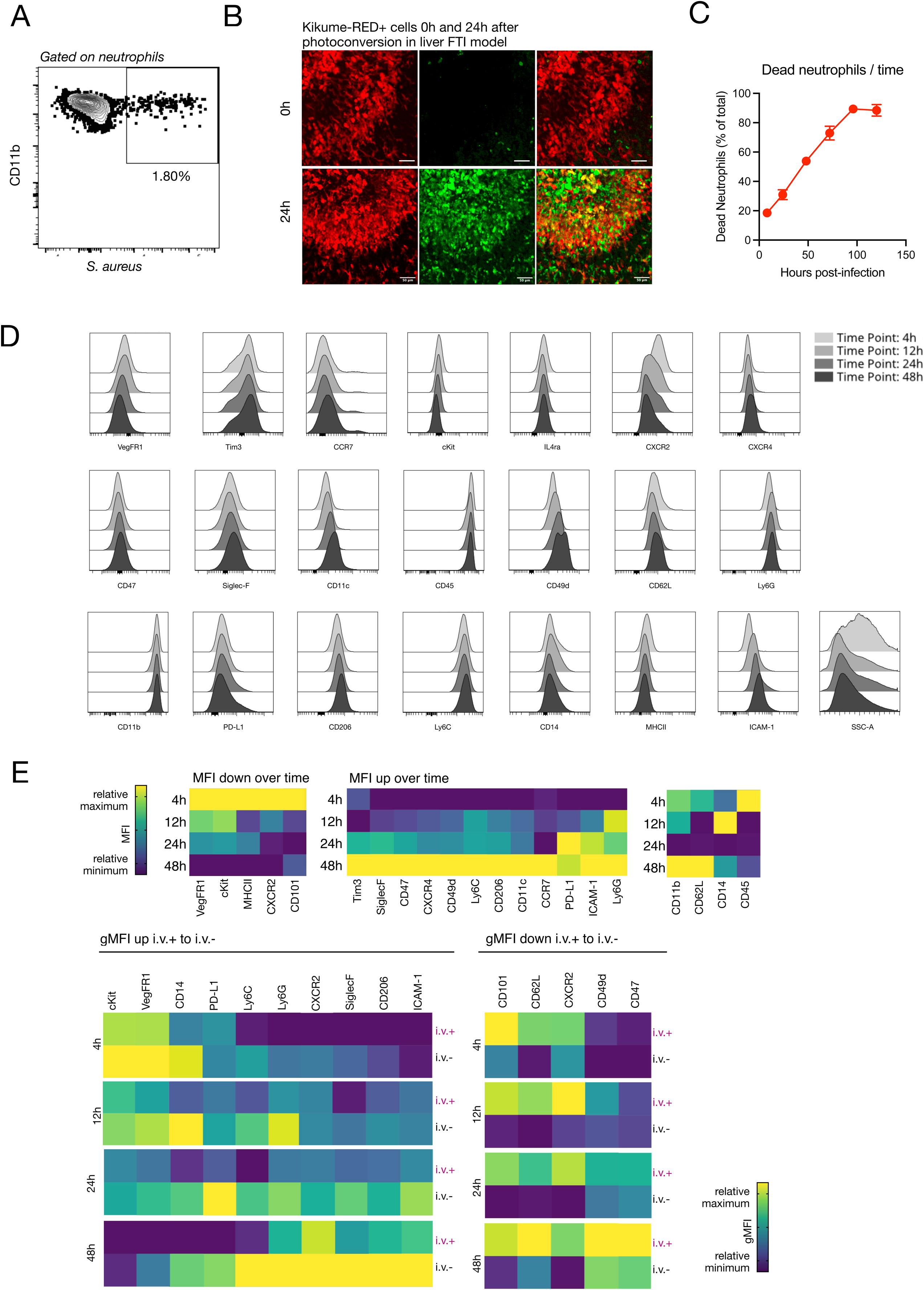
A) Representative flow cytometry data depicting S. aureus positive neutrophils from a skin infection. B) Intravital microscopy images of Kikume-RED+ and Kikume-GREEN+ neutrophils in the liver focal thermal injury model, taken at 0h and 24h after photoactivation. Scale bar = 50 μm C) Quantification of CD11b+ Ly6G+ events within the dead cell gate as a proportion of total CD11b+ Ly6G+ cells (live + dead) over time. Data are taken from multiple experiments and compare values from 8h, 24h, 48h, 72h, 96h, and 120h after infection. N = 4-19 per time point. D) Histogram representations of neutrophil markers across the 4-48h time course. E) Heatmap representations of relative protein expression (fluorescence signal), normalized per each marker, over time and in the iv.v+ versus i.v.- compartments of infections. Error bars = SEM. Related to figure 1.

**Figure S2.**
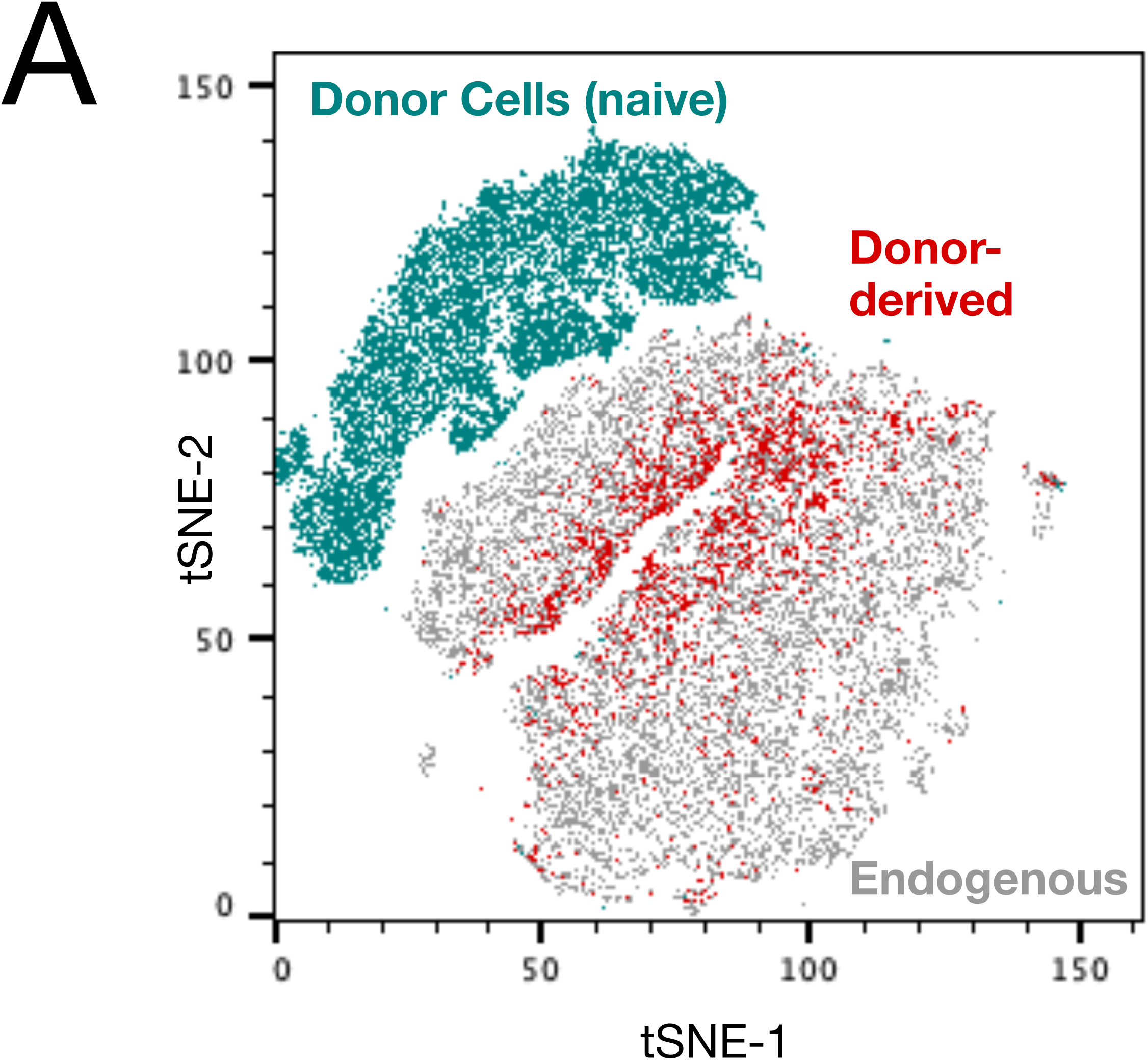
A) tSNE representation of flow cytometry data depicting CD45.1 donor cells before transfer (teal) or 4h after transfer (red) and endogenous cells (gray) from skin infections. N = 3 mice for donor-derived red and gray conditions. Related to figure 2.

**Figure S3.**
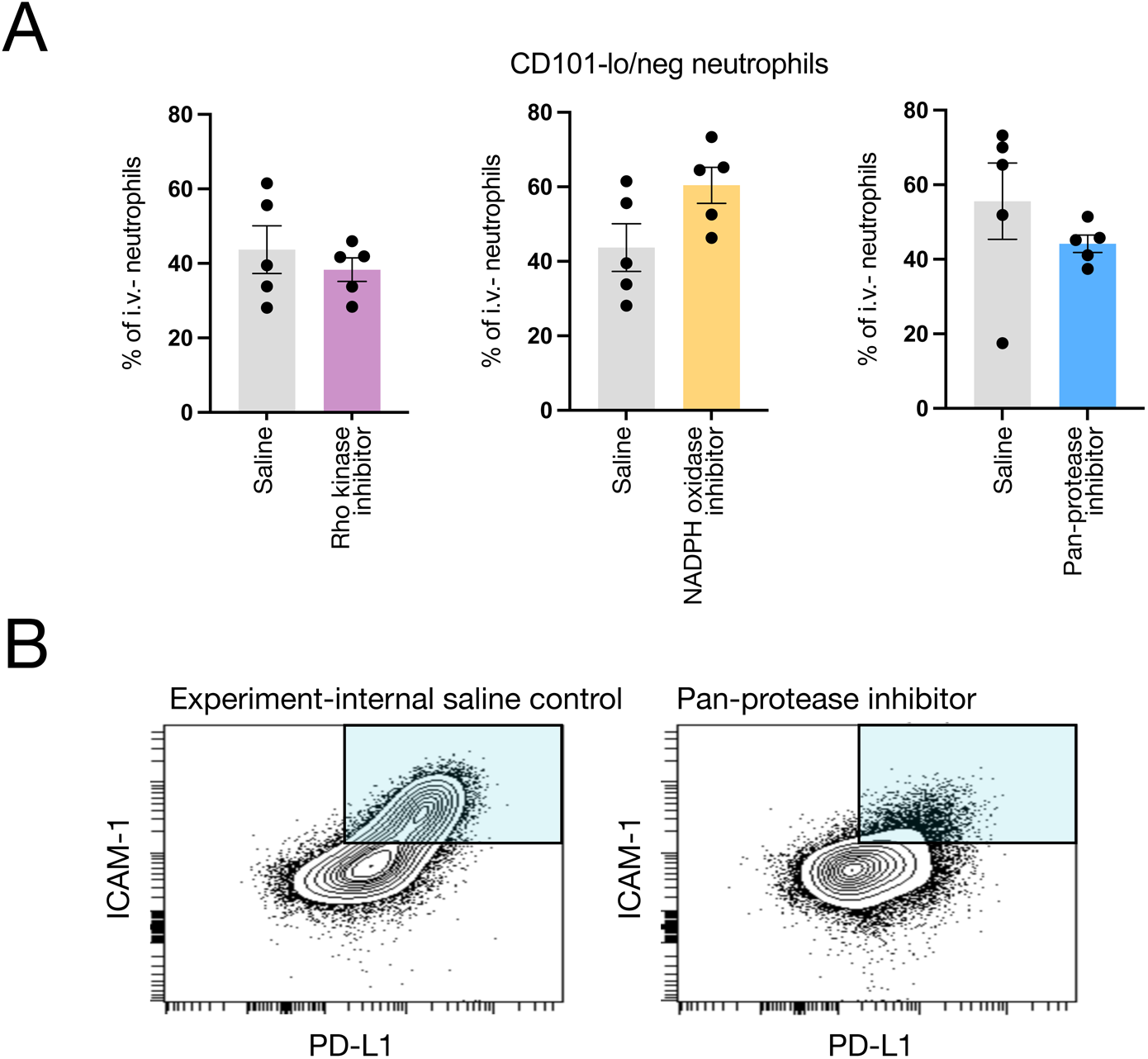
A) Quantification of the proportion of i.v.- neutrophils within the Cd101-lo/negative gate for conditions of NADPH oxidase, Rho kinase, and pan-protease inhibition, compared to saline controls. B) Flow cytometry data depicting the distribution of infection-associated neutrophils in the i.v.- CD101-lo gate per PD-L1 and ICAM-1 expression for pan-protease inhibitor-treated versus experiment-specific saline control. Related to figure 3.

**Figure S4.**
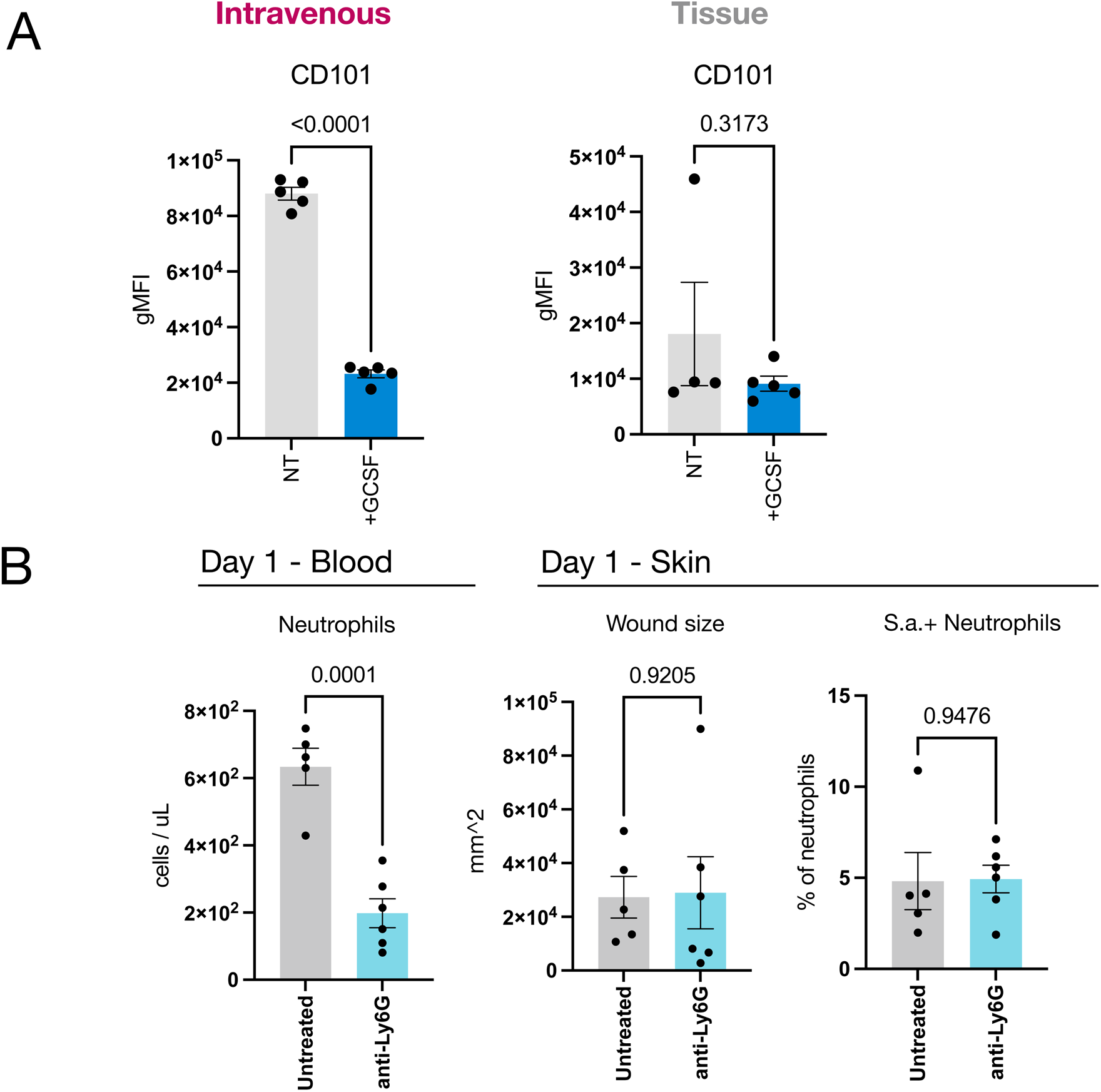
A) Quantification of flow cytometry data depicting neutrophil expression of CD101 in intravenous (left) and tissue-associated (right) neutrophils from control mice or those treated with G-CSF. B) Quantification of flow cytometry data depicting neutrophil levels in blood (left), wound size (middle), and the proportion of *S. aureus*+ neutrophils (right) in 24h infections from mice treated with anti-Ly6G or control mice. N = 5-6 mice per group. Error bars = SEM, statistics = t-test, P values shown. Related to figure 4.

**Figure S5.**
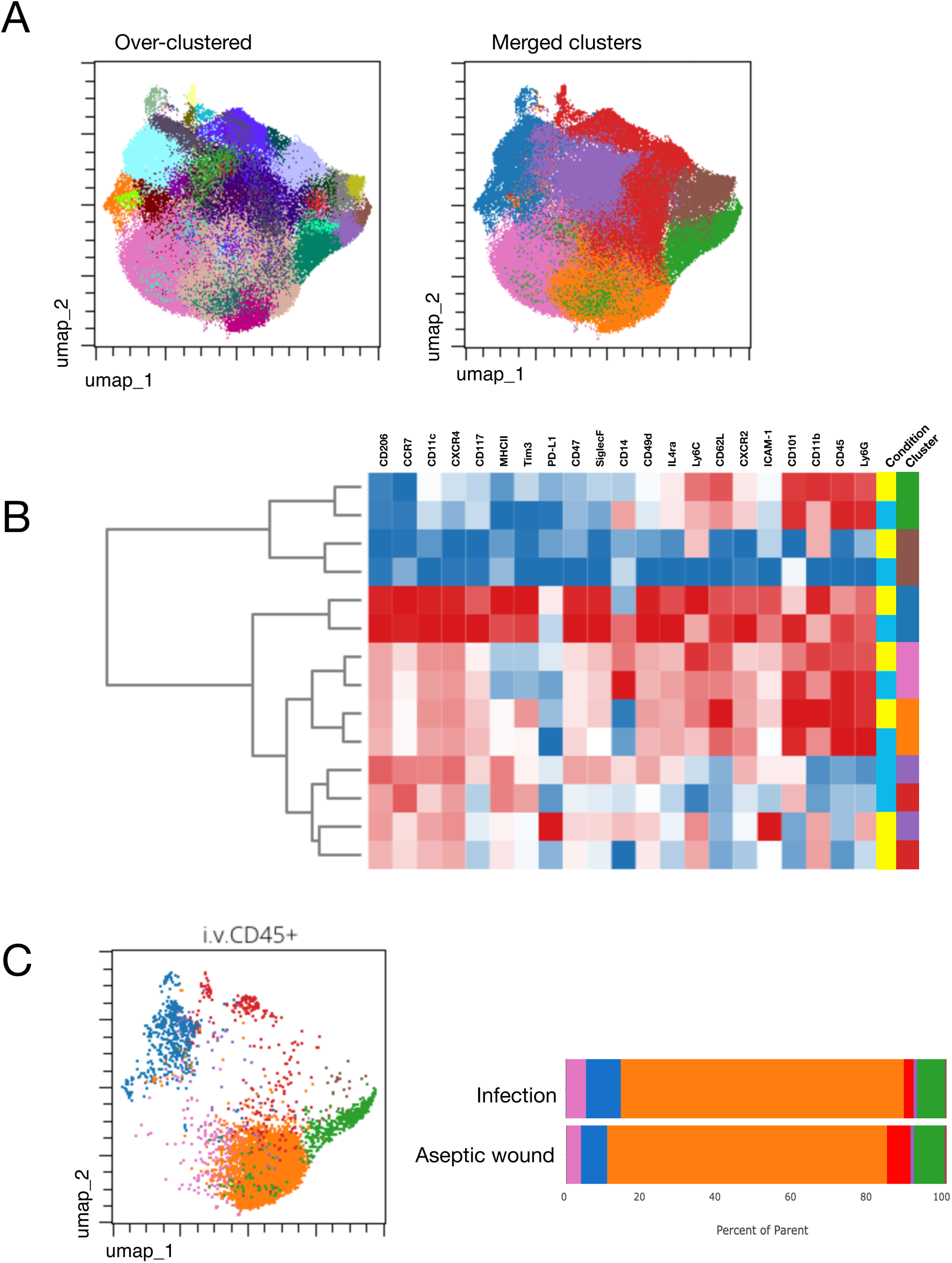
A) Raw output clusters from FlowSOM using comma-separated k values set to 30 (left), and manually merged clusters (right). B) Hierarchically clustered heatmap depicting phenotypic marker expression across clusters, colors scaled per individual maker. C) UMAP representation of i.v. + neutrophils from infections and aseptic wounds (left) and bar plot representation of each cluster as a proportion of total i.v.+ neutrophils for each condition (right). Related to figure 6.

**Figure S6.**
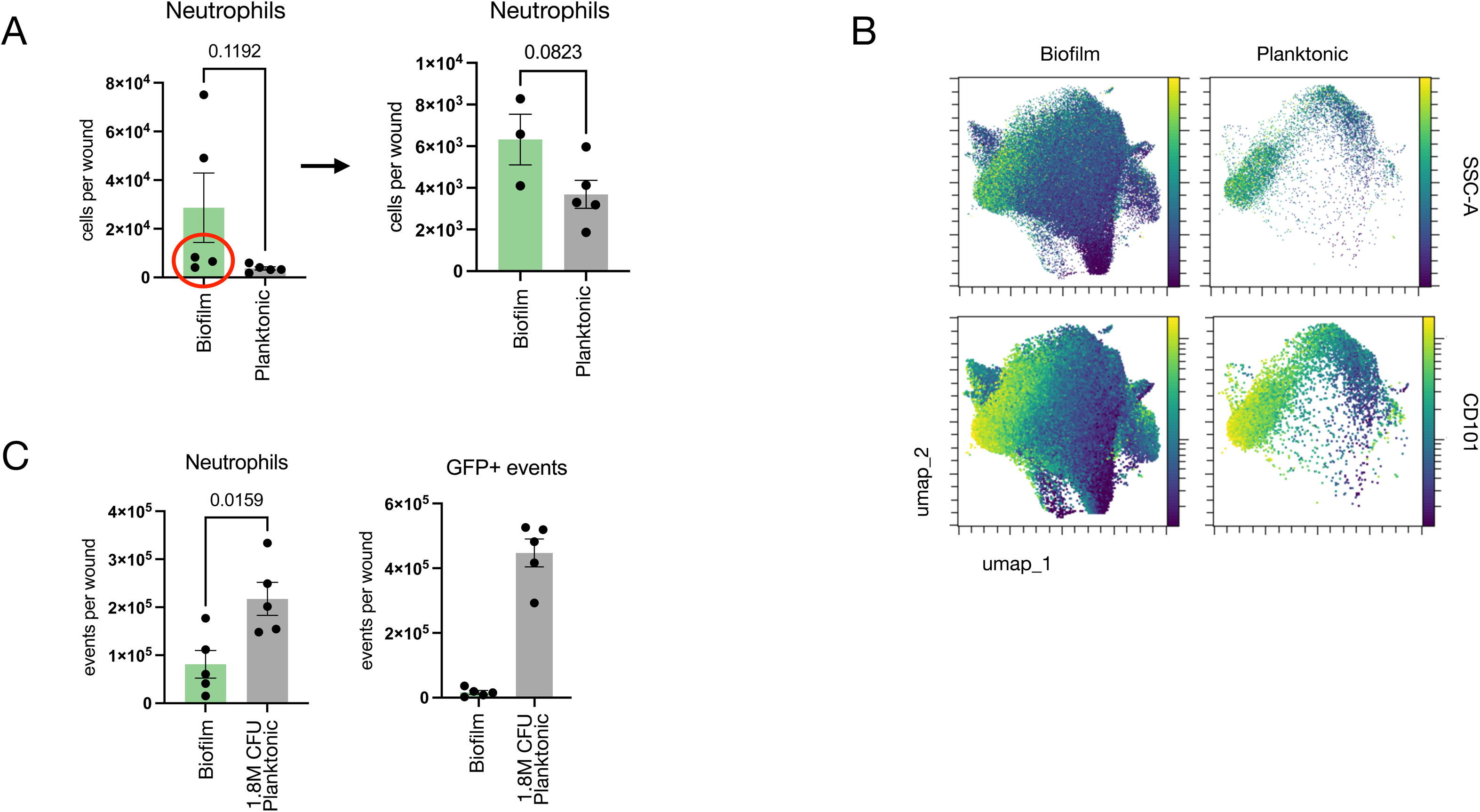
A) Quantification of flow cytometry data depicting numbers of neutrophils in biofilm (left) and planktonic (right) infections, and indicating biological replicates chosen for phenotypic comparisons between biofilm and planktonic infection-associated neutrophils in Figure 7F-H. B) UMAP representations of neutrophils from biofilm (left) and ∼90K CFU MOI planktonic (right) infections, colored by fluorescence intensity reflecting granularity (side scatter, SSC-A, top) or CD101 expression (bottom). C) Quantification of neutrophil numbers (left) and *S. aureus*+ events (right) in biofilm infections versus 1.8M CFU MOI planktonic infections. Error bars = SEM, statistics = t-test, P values shown. Related to figure 7.

## References

1. M. Phillipson, P. Kubes, The Healing Power of Neutrophils Trends Immunol 40, 635–647 (2019).

2. G. Sreejit, J. Johnson, R. M. Jaggers, A. Dahdah, A. J. Murphy, N. M. J. Hanssen, P. R. Nagareddy, Neutrophils in cardiovascular disease: warmongers, peacemakers, or both? Cardiovasc Res 118, 2596–2609 (2022).

3. S. Gupta, M. J. Kaplan, The role of neutrophils and NETosis in autoimmune and renal diseases Nat Rev Nephrol 12, 402–413 (2016).

4. S.-L. Puhl, S. Steffens, Neutrophils in Post-myocardial Infarction Inflammation: Damage vs. Resolution? Front Cardiovasc Med 6, 25 (2019).

5. M. Siwicki, M. J. Pittet, Versatile neutrophil functions in cancer Semin Immunol 57, 101538 (2021).

6. R. Zilionis, C. Engblom, C. Pfirschke, V. Savova, D. Zemmour, H. D. Saatcioglu, I. Krishnan, G. Maroni, C. V. Meyerovitz, C. M. Kerwin, S. Choi, W. G. Richards, A. De Rienzo, D. G. Tenen, R. Bueno, E. Levantini, M. J. Pittet, A. M. Klein, Single-Cell Transcriptomics of Human and Mouse Lung Cancers Reveals Conserved Myeloid Populations across Individuals and Species Immunity 50, 1317–1334.e10 (2019).

7. E. Vafadarnejad, G. Rizzo, L. Krampert, P. Arampatzi, A.-P. Arias-Loza, Y. Nazzal, A. Rizakou, T. Knochenhauer, S. R. Bandi, V. A. Nugroho, D. J. J. Schulz, M. Roesch, P. Alayrac, J. Vilar, J.-S. Silvestre, A. Zernecke, A.-E. Saliba, C. Cochain, Dynamics of Cardiac Neutrophil Diversity in Murine Myocardial Infarction Circ Res 127, e232–e249 (2020).

8. Y. P. Zhu, L. Padgett, H. Q. Dinh, P. Marcovecchio, A. Blatchley, R. Wu, E. Ehinger, C. Kim, Z. Mikulski, G. Seumois, A. Madrigal, P. Vijayanand, C. C. Hedrick, Identification of an Early Unipotent Neutrophil Progenitor with Pro-tumoral Activity in Mouse and Human Bone Marrow Cell Rep 24, 2329–2341.e8 (2018).

9. B. E. Hsu, S. Tabariès, R. M. Johnson, S. Andrzejewski, J. Senecal, C. Lehuédé, M. G. Annis, E. H. Ma, S. Völs, L. Ramsay, R. Froment, A. Monast, I. R. Watson, Z. Granot, R. G. Jones, J. St-Pierre, P. M. Siegel, Immature Low-Density Neutrophils Exhibit Metabolic Flexibility that Facilitates Breast Cancer Liver Metastasis Cell Rep 27, 3902–3915.e6 (2019).

10. C. Pfirschke, C. Engblom, J. Gungabeesoon, Y. Lin, S. Rickelt, R. Zilionis, M. Messemaker, M. Siwicki, G. M. Gerhard, A. Kohl, E. Meylan, R. Weissleder, A. M. Klein, M. J. Pittet, Tumor-Promoting Ly-6G^+^ SiglecF^high^ Cells Are Mature and Long-Lived Neutrophils Cell Rep 32, 108164 (2020).

11. J. Gungabeesoon, N. A. Gort-Freitas, M. Kiss, E. Bolli, M. Messemaker, M. Siwicki, M. Hicham, R. Bill, P. Koch, C. Cianciaruso, F. Duval, C. Pfirschke, M. Mazzola, S. Peters, K. Homicsko, C. Garris, R. Weissleder, A. M. Klein, M. J. Pittet, A neutrophil response linked to tumor control in immunotherapy Cell 186, 1448–1464.e20 (2023).

12. D. M. Calcagno, C. Zhang, A. Toomu, K. Huang, V. K. Ninh, S. Miyamoto, A. D. Aguirre, Z. Fu, J. Heller Brown, K. R. King, SiglecF(HI) Marks Late-Stage Neutrophils of the Infarcted Heart: A Single-Cell Transcriptomic Analysis of Neutrophil Diversification J Am Heart Assoc 10, e019019 (2021).

13. J. Nordvig, T. Aagaard, G. Daugaard, P. Brown, H. Sengeløv, J. Lundgren, M. Helleberg, Febrile Neutropenia and Long-term Risk of Infection Among Patients Treated With Chemotherapy for Malignant Diseases Open Forum Infect Dis 5, ofy255 (2018).

14. L. A. Boxer, Severe congenital neutropenia: genetics and pathogenesis Trans Am Clin Climatol Assoc 117, 13–31; discussion 31 (2006).

15. D. H. McDermott, P. M. Murphy, WHIM syndrome: Immunopathogenesis, treatment and cure strategies Immunol Rev 287, 91–102 (2019).

16. J. Qi, D. D’Souza, T. Dawson, D. Geanon, H. Stefanos, R. Marvin, L. Walker, A. H. Rahman, Multimodal Single-Cell Characterization of the Human Granulocyte Lineage bioRxiv 2021.06.12.448210 (2021).

17. Y. Colino-Sanguino, L. Rodriguez de la Fuente, B. Gloss, A. M. K. Law, K. Handler, M. Pajic, R. Salomon, D. Gallego-Ortega, F. Valdes-Mora, Performance comparison of high throughput single-cell RNA-Seq platforms in complex tissues Heliyon 10, e37185 (2024).

18. P. Fouret, R. M. du Bois, J. F. Bernaudin, H. Takahashi, V. J. Ferrans, R. G. Crystal, Expression of the neutrophil elastase gene during human bone marrow cell differentiation J Exp Med 169, 833–845 (1989).

19. R. M. Kratofil, H. B. Shim, R. Shim, W. Y. Lee, E. Labit, S. Sinha, C. M. Keenan, B. G. J. Surewaard, J. Y. Noh, Y. Sun, K. A. Sharkey, M. Mack, J. Biernaskie, J. F. Deniset, P. Kubes, A monocyte-leptin-angiogenesis pathway critical for repair post-infection Nature 609, 166–173 (2022).

20. S. Nowotschin, A.-K. Hadjantonakis, Use of KikGR a photoconvertible green-to-red fluorescent protein for cell labeling and lineage analysis in ES cells and mouse embryos BMC Dev Biol 9, 49 (2009).

21. M. Evrard, I. W. H. Kwok, S. Z. Chong, K. W. W. Teng, E. Becht, J. Chen, J. L. Sieow, H. L. Penny, G. C. Ching, S. Devi, J. M. Adrover, J. L. Y. Li, K. H. Liong, L. Tan, Z. Poon, S. Foo, J. W. Chua, I.-H. Su, K. Balabanian, F. Bachelerie, S. K. Biswas, A. Larbi, W. Y. K. Hwang, V. Madan, H. P. Koeffler, S. C. Wong, E. W. Newell, A. Hidalgo, F. Ginhoux, L. G. Ng, Developmental Analysis of Bone Marrow Neutrophils Reveals Populations Specialized in Expansion, Trafficking, and Effector Functions Immunity 48, 364–379.e8 (2018).

22. M. S. F. Ng, I. Kwok, L. Tan, C. Shi, D. Cerezo-Wallis, Y. Tan, K. Leong, G. F. Calvo, K. Yang, Y. Zhang, J. Jin, K. H. Liong, D. Wu, R. He, D. Liu, Y. C. Teh, C. Bleriot, N. Caronni, Z. Liu, K. Duan, V. Narang, I. Ballesteros, F. Moalli, M. Li, J. Chen, Y. Liu, L. Liu, J. Qi, Y. Liu, L. Jiang, B. Shen, H. Cheng, T. Cheng, V. Angeli, A. Sharma, Y.-H. Loh, H. L. Tey, S. Z. Chong, M. Iannacone, R. Ostuni, A. Hidalgo, F. Ginhoux, L. G. Ng, Deterministic reprogramming of neutrophils within tumors Science 383, eadf6493 (2024).

23. G. Boivin, J. Faget, P.-B. Ancey, A. Gkasti, J. Mussard, C. Engblom, C. Pfirschke, C. Contat, J. Pascual, J. Vazquez, N. Bendriss-Vermare, C. Caux, M.-C. Vozenin, M. J. Pittet, M. Gunzer, E. Meylan, Durable and controlled depletion of neutrophils in mice Nat Commun 11, 2762 (2020).

24. K. Moses, J. C. Klein, L. Männ, A. Klingberg, M. Gunzer, S. Brandau, Survival of residual neutrophils and accelerated myelopoiesis limit the efficacy of antibody-mediated depletion of Ly-6G+ cells in tumor-bearing mice J Leukoc Biol 99, 811–823 (2016).

25. J. P. Mohammed, M. E. Fusakio, D. B. Rainbow, C. Moule, H. I. Fraser, J. Clark, J. A. Todd, L. B. Peterson, P. B. Savage, M. Wills-Karp, W. M. Ridgway, L. S. Wicker, J. Mattner, Identification of Cd101 as a Susceptibility Gene for Novosphingobium aromaticivorans-Induced Liver Autoimmunity The Journal of Immunology J Immunol 187, 337–349 (2011).

